# Epistatic interactions shape the interplay between beneficial alleles and gain or loss of pathways in the evolution of novel metabolism

**DOI:** 10.1101/2020.10.20.347948

**Authors:** Eric L. Bruger, Lon M. Chubiz, José I. Rojas Echenique, Caleb J. Renshaw, N. Victoria Espericueta, Jeremy A. Draghi, Christopher J. Marx

## Abstract

Fitness landscapes are often invoked to interpret the effects of allele substitutions and their interactions; however, evolution also includes larger changes like gene loss and acquisition. Previous work with the methylotrophic bacterium *Methylorubrum extorquens* AM1 identified strongly beneficial mutations in a strain evolved to utilize a novel, *Foreign* pathway in place of its native central metabolic pathway for growth on methanol. These mutations were consistently beneficial, regardless of the order in which they arose. Here we extend this analysis to consider loss or acquisition of metabolic pathways by examining strains relying upon either the *Native* pathway, or both (‘*Dual*’) pathways present. Unlike in the *Foreign* pathway context in which they evolved, these alleles were often deleterious in these alternative genetic backgrounds, following patterns that were strongly contingent on the specific pathways and other evolved alleles present. Landscapes for these alternative pathway backgrounds altered which genotypes correspond to local fitness peaks and would restrict the set of accessible evolutionary trajectories. These epistatic interactions negatively impact the probability of maintaining multiple degenerate pathways, making it more difficult for these pathways to coevolve. Together, our results highlight the uncertainty of retaining novel functions acquired via horizontal gene transfer (HGT), and that the potential for cells to either adopt novel functions or to maintain degenerate pathways together in a genome is heavily dependent upon the underlying epistatic interactions between them.

**Author Summary:** The evolution of physiology in microbes has important impacts ranging from global cycling of elements to the emergence and spread of pathogens and their resistance to antibiotics. While genetic interactions between mutations in evolving lineages of microbes have been investigated, these have not included the acquisition of novel genes on elements like plasmids, and thus how these elements interact with existing alleles. The dynamics of novel gene retention are of interest from both positive (e.g., biotechnology) and negative (e.g., antimicrobial resistance) practical impacts. We find that the patterns of interactions between evolved alleles appear substantially different, and generally much less positive, when moved into novel genetic backgrounds. Additionally, these preexisting alleles were found to have strong impacts on the ability of genotypes to maintain – and in rare cases coevolve with – novel genes and pathways. These results show that even though they evolved separately, the particular alleles in a genetic background, and importantly the physiological impacts they confer, weigh heavily on whether genes for novel metabolic processes are maintained.

## Introduction

The means by which novel functions like metabolic pathways evolve is an open and highly relevant question for all biological systems [1,2]. One existing path to novelty is the evolutionary diversification of functionally analogous pathways from a common ancestral function [3–6]. Alternatively, novel functions can be obtained via horizontal gene transfer (HGT). The genetic bases of these novel functions could be paralogs of an existing pathway, thus providing redundancy, or pathways that execute the same function or yield similar outputs/products with unrelated, structurally different components, and thus conferring degeneracy [7–10]. Although degeneracy is a prominent feature of biological systems such as cellular metabolism, the evolutionary routes to degeneracy at the systems level (e.g., in metabolic networks) are not well understood. This study aims to address a question fundamental to the evolution of metabolism: what drives the maintenance of – and sometimes replacement – of ancestral pathways by foreign, non-homologous pathways (Fig. 1)?

**Figure 1.**
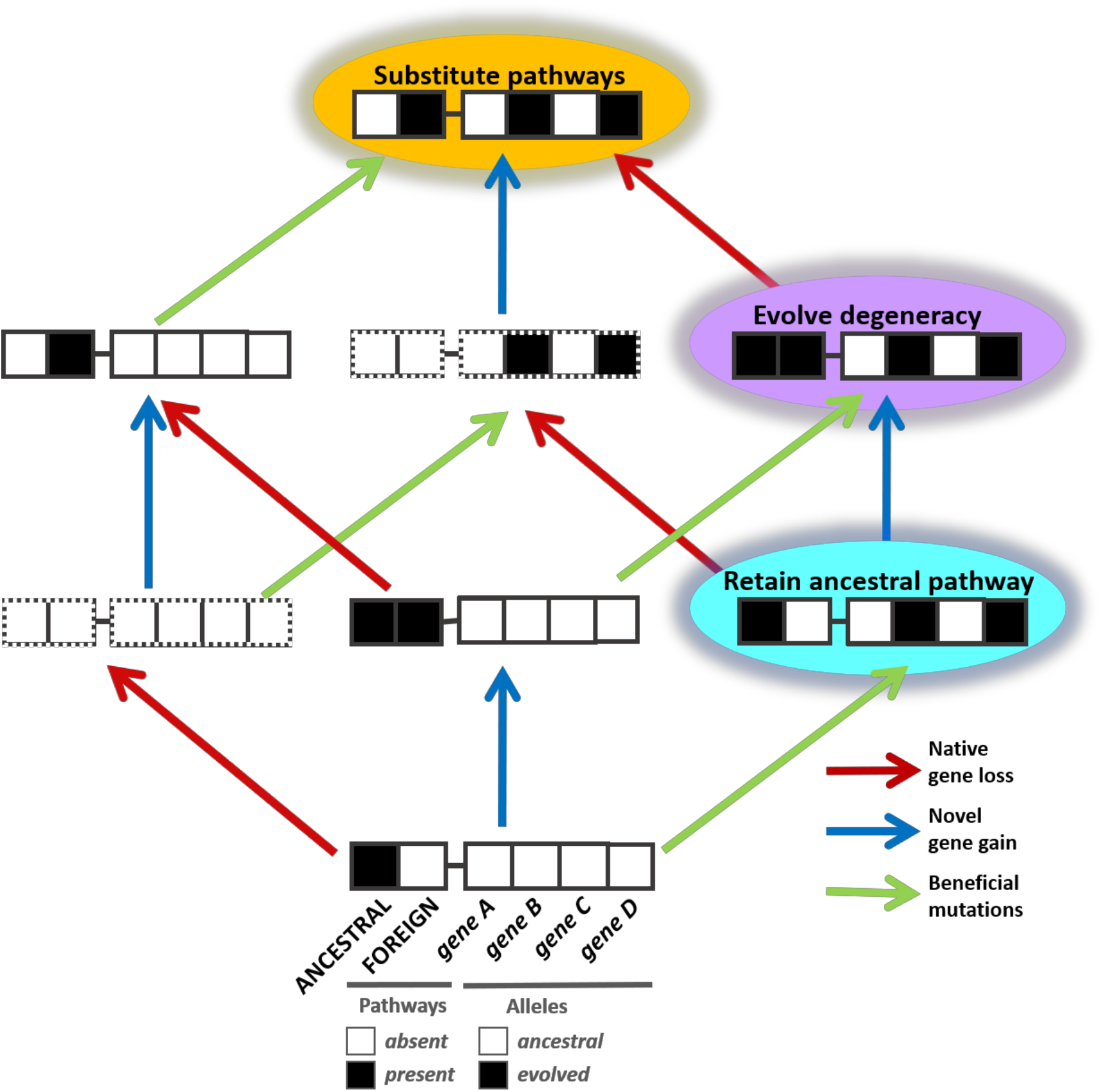
Combined impacts of metabolic gene gain, loss, and beneficial mutation on competitive fitness. For a given process, required and redundant genes can be gained and loss, and additional mutations that improve fitness can be acquired and selected. Epistasis, both in terms of sign and magnitude, can emerge between ancestral, evolved, and newly acquired alleles. Presence of the ancestral pathway is indicated by a red box, inheriting a novel pathway is marked by a blue box, and a green box represents the evolution of beneficial alleles. Boxes bordered by broken lines indicate genotypes that are nonviable under the selective condition examined, due to the absence of any metabolic pathway for the required metabolic process. Abbreviations are as follows: “ANC” = ancestral pathway, “FOR” = novel foreign pathway, “EVOL” = acquisition of evolved beneficial alleles.

This study examines how genetic context shapes the evolution of metabolic pathways using methylotrophy as a model system (Fig. 2). Methylotrophic organisms have the unique metabolic ability to utilize one-carbon compounds as sole sources of carbon and energy. A direct consequence of methylotrophic metabolism is that all carbon flows through the toxic intermediate, formaldehyde. Four known pathways for formaldehyde oxidation exist in bacteria [11]. The functional redundancy provided by degeneracy is a common feature among methylotrophs, as many methylotrophs harbor more than one of these pathways in tandem. It has been hypothesized that selection has maintained these degenerate pathways by providing multiple routes to prevent formaldehyde accumulation [11–13]. The model methylotroph *Methylorubrum extorquens* AM1 encodes a single formaldehyde oxidation pathway that uses the cofactor dephospho-tetrahydromethanopterin (dH_4_MPT; referred to throughout as the ‘*Native*’ or dH_4_MPT pathway; Fig. 2) [14].

**Figure 2.**
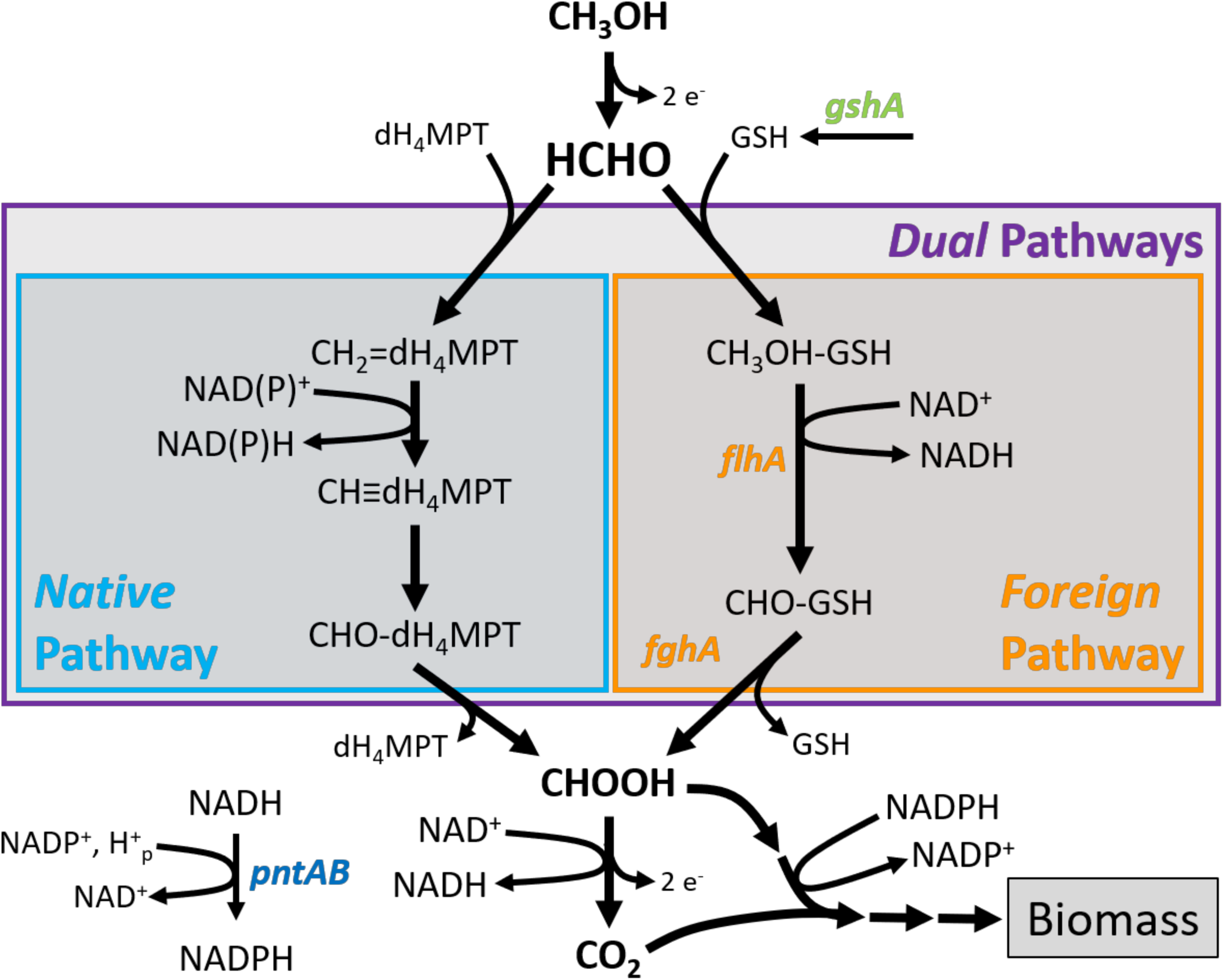
Formaldehyde oxidation pathways used in this study. The “*Native*” pathway for formaldehyde oxidation normally found in *Methylorubrum extorquens* AM1 is the dH_4_MPT-dependent branch displayed. Alternatively, the “*Foreign*” degenerate GSH-dependent pathway from *Paracoccus denitrificans* was cloned onto plasmid pCM410 and moved into *M. extorquens*. “*Dual*” pathway strains have degeneracy, as they possess both of these unrelated pathways.

During the process of determining the role of the dH_4_MPT pathway in *M. extorquens*, it was found that this *Native* pathway could be replaced with a distinct, unrelated formaldehyde oxidation pathway from another bacterium [15]. Two enzymes of the glutathione (GSH)-dependent formaldehyde oxidation pathway from *Paracoccus denitrificans* [16] were introduced into a *M. extorquens* strain in which the dH_4_MPT pathway had been disrupted by a deletion of *mptG*, encoding β-ribofuranosylaminobenzene 5’-phosphate synthase [15]. This allowed the ‘*Foreign*’ GSH pathway to serve as a substitute pathway for formaldehyde oxidation to formate (Fig. 2). This pathway is composed of *S*-hydroxymethyl-GSH dehydrogenase [16] and *S*-formyl-GSH hydrolase [17], encoded by *flhA* and *fghA*, respectively. Although functionally analogous to the *Native* dH_4_MPT pathway in terms of converting formaldehyde to formate, the enzymes of the *Foreign* GSH pathway can only couple to NAD^+^, generating NADH. In contrast, the analogous enzymes of the dH_4_MPT pathway can couple to either NAD^+^ or NADP^+^, thereby generating NADH or NADPH [14]. Since this step is the only central metabolic reaction generating NADPH during growth on methanol, the exclusive use of the *Foreign* pathway also requires involvement of transhydrogenase (encoded by *pntAB*) to convert NADH (and NADP^+^) to NADPH (and NAD^+^) while consuming one proton from the proton motive force in the process [18].

The *M. extorquens* strain with a *Foreign* formaldehyde oxidation pathway replacing its *Native* one could grow on methanol, but because it grew approximately a threefold worse than wild-type, experimental evolution was used to select for improved growth [19]. These populations rapidly re-evolved faster growth, and an isolate from a particularly fit population was sequenced to identify candidate beneficial mutations in that strain [19]. Beneficial mutations were identified in: A) *fghA/flhA*, alleviating the gene expression burden [20,21]; B) *pntAB*, increasing expression of the key step that generates NADPH for biosynthetic reactions [22]; C) *gshA* (encoding γ-glutamylcysteine synthetase), which likely results in increased cellular GSH pools; and D) six other loci without obvious connection to formaldehyde oxidation (which was treated as a single ‘allele’ designated as ‘*GB*’ for ‘Genetic Background’) [19]. Beneficial mutations in *fghA*, *pntAB*, and *gshA* were identified across multiple populations [20,22], thus they were repeated targets for improvement. These alleles evolved in the strain that was sequenced in the following order: *gshA*, *fghA*, then *pntAB* [23]. The fitness gains of the strains evolved with the *Foreign* pathway were found to be specific to growth on methanol or methylamine, two C_1_ substrates that generate formaldehyde, rather than growth on the multi-C substrate succinate, nor the C_1_ substrate formate that is downstream of formaldehyde oxidation [19,22]. This suggested that adaptation had been fairly specific to improving the ability of cells to use the *Foreign* GSH pathway as a central metabolic pathway.

Are all adaptive trajectories leading to this evolved genotype possible and likely, or do interactions among mutant alleles constrain their likely sequence of evolution? This was addressed by generating all potential genotypic intermediates (2^4^ = 16 combinations) of these four beneficial alleles [19] in order to identify epistatic interactions that might exist between them. Epistatic, or non-linear, interactions between alleles occur when mutations have different effects on fitness (or other traits) in combination than expected by comparing their effects in isolation [24,25]. This can lead to differences in either the magnitude and/or sign of selection [19,26–35]. These diverse patterns of epistasis affect other system features including robustness, evolvability, genome complexity, and genetic architecture in complex ways that are not fully understood [9,36–41]. In the context of the fitness landscape metaphor [42–46], epistasis produces the curvature and ruggedness in the genotype-fitness terrain. The terrain of the landscape indicates whether a given evolutionary path will be accessible, defined as adaptive at each step along the sequence of substitutions. In the metabolic system originally examined in this study, it was found that all derived alleles were beneficial on all backgrounds (i.e., no sign epistasis). This finding suggested that the strain bearing all evolved alleles represents the fitness peak among the possible combinatorial genotypes and that the landscape is fairly smooth with no local maxima or fitness valleys to cross [47–50]. Conversely, where there is sign epistasis – alleles being beneficial on some backgrounds and deleterious on others – some adaptive trajectories become selectively inaccessible because they require transitions across a fitness valley [26,51–53].

Would the same changes that render a novel pathway more fit also facilitate the acquisition of multiple, degenerate pathways, or would they instead interfere with a strain’s ability to utilize its original cellular machinery? This question hinges entirely upon the shape of epistatic interactions. At one level are the interactions between the new beneficial mutations that emerged in the novel context of a *Foreign* pathway and other contexts, which could be thought of as “*mutation × pathway*” interactions. At a second level, the pathway context can also influence the manner in which beneficial mutations interact with each other, or “*mutation × mutation × pathway*” interaction. As stated above, in the original context of the evolution of strains with the *Foreign* GSH pathway in place of the *Native* dH_4_MPT pathway all mutations were individually beneficial. What might be expected for these mutations in genotypes that contain the *Native* pathway, alone or in the *Dual* context? Mutations that improve a generic function needed for growth on methanol, or even simply the lab conditions, would be expected to remain beneficial. But given that use of the GSH pathway presents unique challenges, it might also be hypothesized that these would either be irrelevant to strains with the dH_4_MPT pathway (i.e., neutral) or generate tradeoffs between the use of one pathway versus the other (i.e., deleterious). What might be expected for the “*mutation × mutation × pathway*” interactions? Although all beneficial mutations were found to be beneficial on the other genotypes in the *Foreign* pathway context they evolved, this may no longer be true. If one or more of these mutations generate tradeoffs between pathway use, this may lead to changes in the sign of their effects upon some backgrounds when the *Native* pathway is present.

Perhaps the most important epistatic interaction in considering the evolution of degeneracy is whether maintaining the novel function is beneficial, neutral, or deleterious. If the second pathway is immediately an advantage, and maintenance of the original pathway remains beneficial, the emergence of degeneracy is straightforward. In contrast, if the novel pathway is initially neutral or deleterious in the presence of the endogenous pathway, then loss will occur unless adaptation towards degeneracy can occur rapidly enough. The *Foreign* GSH pathway was shown to be costly in WT *M. extorquens* that has its *Native* dH_4_MPT pathway due to both plasmid maintenance and enzyme expression costs [20,54], but it remains unclear whether these costs might be reduced or even eliminated in the presence of other beneficial alleles that improved its function in isolation.

Here we take a first step to considering adaptation against the backdrop of the gain or loss of metabolic pathways by asking how mutations that evolved in the context of the *Foreign* pathway would act in strains that carry either the *Native* pathway alone or the *Dual* pathway strain. To date, we are unaware of any other studies that examine the impact of degeneracy upon epistasis and fitness landscapes by integrating examination of genetic changes available via mutation (alternative alleles or gene loss) with those involving gene gain through HGT. This approach allows us to address several questions pertaining to the evolution of degeneracy and/or metabolic novelty: 1) Do patterns of epistasis between alleles that evolved in the presence of a particular pathway change in the presence of alternative pathway(s)?; 2) Do these changes impact which evolutionary trajectories are accessible?; 3) How do these changes impact the ability of cells to retain and evolve with multiple pathways for a given metabolic process? Whereas sign epistasis is absent in the context of the *Foreign* pathway [19], we found it was prevalent in both the *Dual* and *Native* contexts. These deleterious effects introduce inaccessible trajectories that would constrain the evolutionary routes and end-points among combinations of these alleles [19,23,55]. The *Dual* strains face a particular challenge, as the presence of the *Foreign* pathway was found to be quite costly initially. However, compared to their respective ancestors, several *Dual* strains with alleles that arose during evolution of the *Foreign* pathway alone had increased plasmid retention, including one case of a *Dual* strain maintaining the GSH pathway and adapting fast enough to remain common in the population after ~100 generations. This highlights the potential impact of epistatic interactions between pathways and evolved alleles on whether cells retain and evolve to use newly acquired metabolic pathways, either alone or in combination with preexisting metabolic functions.

## Methods

### Bacterial strains

A full list of the *M. extorquens* strains used in this study is described in Table S1. The strains under analysis for epistatic comparisons contain combinations of evolved alleles previous described, categorized as *fghA*, *pntAB*, *gshA*, and *GB* mutations [19]. The Genetic Background (‘GB’) allele is actually a composite of six additional mutations that occurred in the CM1145 population (1 SNP on plasmid, loss of a plasmid, 2 IS insertions, 1 small insertion, and 1 large deletion) [19]. One of these IS insertions [56] and the large deletion [21] were commonly observed in populations of WT *M. extorquens* evolving in other nutritional environments and were determined to be beneficial for traits unlinked to formaldehyde oxidation. None of the other four loci have an obvious connection to methylotrophy. For convenience, the genotypes used are often referred to by their bitstring representations at the focal alleles under examination, where an allelic state of “0” represents the ancestral (“^anc^”) allele, a “1” represents the evolved (“^evo^”), an a “-“ represents the absence of an allele/pathway (Fig. S1A, Table S1). Presence or absence of a given pathway are indicated by “1” and “0”, respectively

### Media and chemicals

All growth experiments were carried out in modified Hypho growth media [56–58]. One liter of growth media consisted of 1 mL of a trace metal solution (12.738 g of EDTA disodium salt dihydrate, 4.4 g of ZnSO_4_· 7H_2_O, 1.466 g of CaCl_2_· 2H_2_O, 1.012 g of MnCl_2_·4H_2_O, 0.22 g of (NH_4_)_6_M_o7_O_24_· 4H_2_O, 0.314 g of CuSO_4_· 5H_2_O, 0.322 g of CoCl_2_·6H_2_O, and 0.998 g of FeSO_4_·7H_2_O per liter), 100 mL of phosphate buffer (25.3 g of K_2_HPO_4_ and 22.5 g of NaH_2_PO_4_ in 1 liter of deionized water), 100 mL of sulfate solution (5 g of (NH_4_)_2_SO_4_ and 0.98 g of MgSO_4_ in 1 liter of deionized water), 799 mL of deionized water, and the desired carbon source. Carbon was supplemented to the media as disodium succinate (3.5 mM, Fisher) or anhydrous methanol (15 mM, Macron Fine Chemicals). The modified trace metal mix (per 100 mL) consisted of 10 mL of 179.5 mM FeSO_4_ (5x relative to above recipe), 80 mL of premixed metal mix (12.738 g of EDTA disodium salt dihydrate, 4.4 g of ZnSO_4_· 7H_2_O, 1.466 g of CaCl_2_· 2H_2_O, 1.012 g of MnCl_2_· 4H_2_O, 0.22 g of (NH_4_)_6_M_o7_O_24_· 4H_2_O, 0.314 g of CuSO_4_· 5H_2_O, and 0.322 g of CoCl_2_· 6H_2_O in 1 liter of deionized water, pH 5), and 10 mL of deionized water.

### Competition Fitness Assays

All cultures were grown at 30 °C in 5 mL of media in sealed Balch tubes (22 mL total capacity) that were aerated using roller drums rotating at approximately 60 rpm. Starter cultures for all strains were initiated from single colonies into Hypho media containing 7.5 mM methanol and 1.75 mM succinate for 2 days. The cultures were then diluted 1/64 into acclimation cultures of Hypho media containing 15 mM methanol for 2-4 days (depending on the growth rate of a given strain in methanol). Finally, cultures were mixed and diluted 1/64 into homologous Hypho media containing 15 mM methanol and competed for a duration of 4 days. All competition assays were initiated as a 1:1 volumetric ratio against a *mCherry* fluorescently-labeled reference strain (CM1232 or CM1176, see Table S1) as previously [20]. Frequencies of competitors were determined by passing mixed population samples from the start (F_0_) and end (F_1_) of the experiment through a Cytoflex S flow cytometer (Beckman Coulter, Indianapolis, Indiana). Approximately 50,000 cells per sample were gated by forward and side scatter and competitors were differentiated through comparison of fluorescence by excitation at 561 nm and emission through a 610/20 BP nm filter. Where required, additional control strains chromosomally tagged with the Venus fluorescent reporter gene were also included and assessed by excitation at 488 nm and detection of emission with a 525/40 BP nm filter. Mathusian fitness measurements (*W*) were determined by ratios of log-transformed changes in relative frequencies over the course of the experiment relative to a common competitor strain (ancestor CM1232, or wild-type CM1176), by the following formula that assumes an average of 64-fold (2^6^) expanded size of mixed populations during competition experiments [19,59]: *W* = log(F_1_*64/F_0_)/log((1-F_1_)*64/(1-F_0_)).

To qualify the conditions used in this study, we compared these results against conditions used previously. The correlation between the fitness values reported previously with cells grown in 50 mL flasks and the new data with cultures grown in tubes placed in a roller drum due to the large quantity involved was high (R^2^ = 0.9, Fig. S2A). Directly comparing new and old strains both grown in roller drums indicated that it was growth conditions and/or the flow cytometry-based fitness assay that generated the remaining differences, not the strains themselves (Fig. S2B). One reason that this correlation has a slope below unity is due to a confounding effect of cell size and distinguishing fluorescent from nonfluorescent strains via flow cytometry that was discovered in subsequent work that led to a consistent underestimate of fitness values for these strains in the initial 2011 study [19]. Because the *Native* and *Dual* sets of strains had the *Native* dH_4_MPT pathway, they were much fitter than the *Foreign* pathway strains, even the fittest strain in that context which had all four evolved alleles present (Fig. 3). Because of this, a fluorescent version of the WT strain (10-000) was used for competition assays to provide more precise fitness estimates and the correlation between both competitor strain approaches was reasonably high (R^2^ = 0.68, Fig. S3). However, for dH_4_MPT pathway-containing strains, we opted to use the data from competitions against wild-type because they allowed better resolution between strains and this data had lower levels of variation on the whole, largely as a result of dH_4_MPT pathway strains having much higher fitness values in comparison to CM1232 (average coefficient of variation of 1.9% vs. 4.7%).

**Figure 3.**
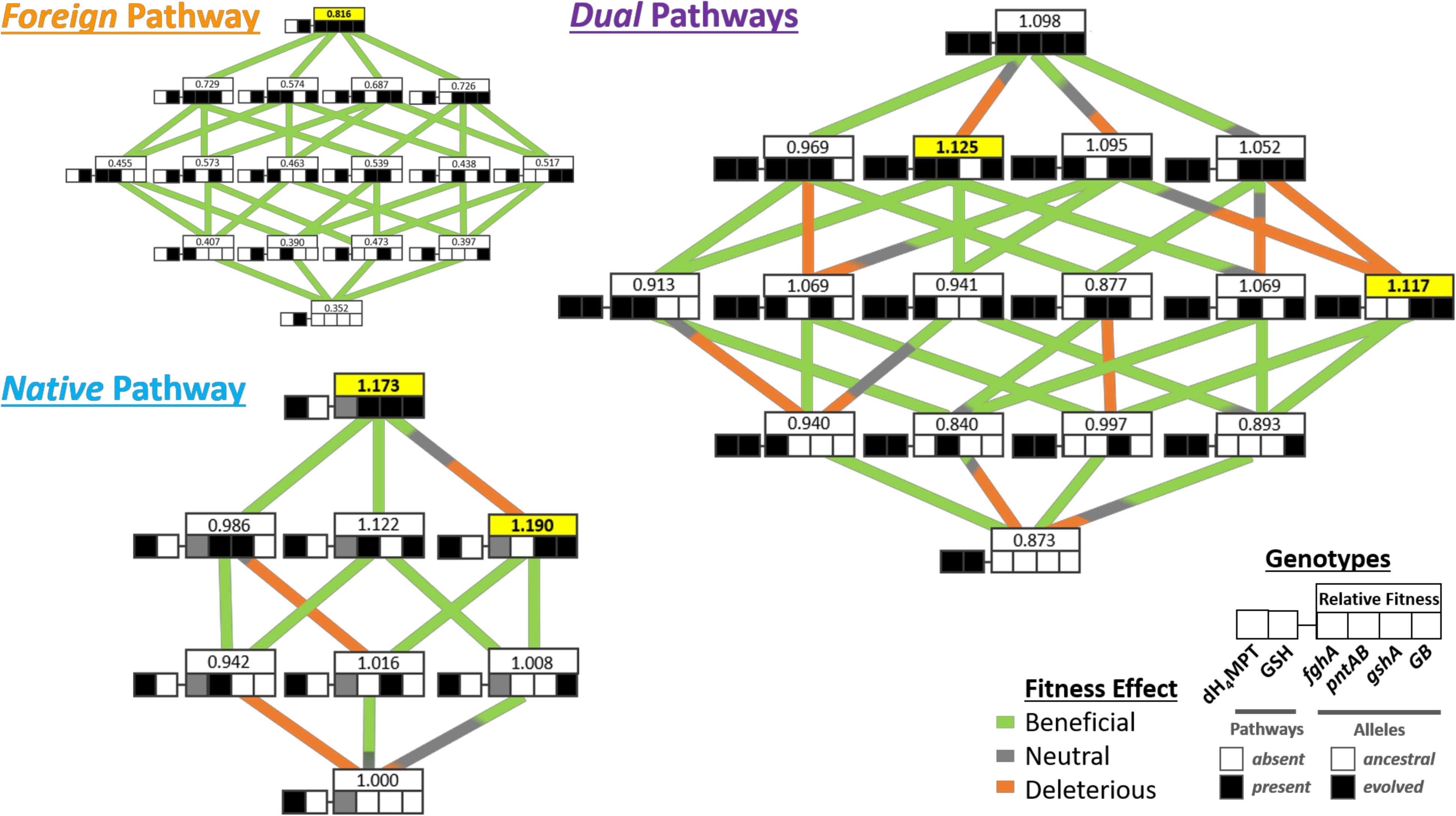
Fitness landscape for genotypes of all pathway backgrounds. **A)** All *Foreign* strains were competed pairwise with the reference strain CM1232 (genotype ’01-0000‘) B) *Dual* and C) *Native* strains were competed pairwise with the reference strain CM1176 (genotype ’10-x000’). For more information on strains see Table S1. Three major subsections of the landscape exist: A) the *Foreign* pathway strains that contain only the GSH pathway, B) the *Dual* pathway strains that contain both the *Native* pathway and the *Foreign* GSH pathway, and C) *Native* pathway strains that contain only the dH_4_MPT pathway. In the diagram, each row of boxes depicts a single strain genotype, with the genotype of the focal alleles represented by the coloration pattern of the boxes. The leftmost box represents the Pathway background of a given genotype and the right four boxes depict the four focal alleles (from left to right: *fghA*, *pntAB*, *gshA*, and *GB*) in either their ancestral (white) or evolved (black) states. The *fghA* box for the *Native* pathway strains is grayed out because the allele is absent in these genotypes. The lines connecting genotypes represent the transitions between mutational neighbors and are color-coded by their net effect: beneficial in green, deleterious in orange, and neutral in gray. Additionally, line thicknesses are weighted by the probabilities of the mutational changes from simulated fitness draws.

In order to confirm that our fitness measurements were broadly consistent with prior work, we compared our results to the *Foreign* combinations previously examined [19]; these strains were regenerated so that all strains considered here shared an identical genetic platform. As reported previously, all permutations of evolved alleles were found to be beneficial in the context of the *Foreign* pathway alone (Fig. 3A, Tables S2 & S3). The ordering of selective coefficients of the individual alleles (*gshA*^evo^, *fghA*^evo^, *GB*^evo^, *pntAB*^evo^) was likewise maintained (Fig. 3A, Table S2 & S3). The largest difference between the data sets was that combinations with 3 or more evolved alleles were found to outperform previous observations. Despite this, the overall landscape remained smooth and universally accessible (Figs. S2 & 3).

### Statistical Comparisons

Model fits were made with linear regression in R (version 3.6.1). Analysis of growth (Fig. S5) was conducted using GraphPad Prism 8.4.3. Analysis of transitions and trajectories over fitness landscapes were conducted by simulating random draws of relative fitness values from the fitness distributions of all genotypes in the landscape over 10^6^ replicate draws (Fig. S6). Fitness distributions were assumed to be Gaussian, parametrized by the sample mean and variances. Among the *Native* and *Dual* strains, alleles that had slightly negative selective coefficients on average were still likely to generate beneficial draws a significant portion of the time due to uncertainty in the fitness measurements (Fig. S4). Probabilities of transitions or trajectories being beneficial were then calculated as the fraction of draws in this ensemble for which a given quantitative relationship (e.g., *S*_evolved_ > 1%) is true. Individual transitions were assigned a designation as beneficial, deleterious, or neutral based on the net probabilities that draws fell into these different classes (Fig. S1B). The probability of reaching the genotype in each particular pathway background with evolved alleles at each locus is the proportion of draws in which all steps along that trajectory had *S*_evolved_ > 1%. Where reported, selective coefficients (*S*) are calculated as previously under a multiplicative model [60,61] as *S* = *W*_evolved_/*W*_ancestor_ – 1.

### Growth assays

To assess growth properties of a subset of strains in the study (Fig. S5), cell cultures were initiated identically to competition experiments, going through starter and acclimation cultures prior to actual growth experiments. Cultures were grown in identical Hypho media in sealed Balch tubes (5 mL media in 22 mL total capacity), and growth was monitored by measurement in a Spectronic 200 spectrophotometer.

### Genetic techniques

Evolved alleles identified in previous studies [19] were combined in all possible permutations into *Native* (wild-type) and *Dual* (wild-type + plasmid) pathway strains. Chromosomal mutations were introduced by allelic exchange as previously described [62] and plasmid-based mutations were introduced by transformation of plasmid variants (i.e., pCM410 and the evolved derivative pCM410.1145, [19]).

### Plasmid carriage experiments

Biological triplicate samples of 5 genotypes from the *Dual* pathway strain set (11-0000, 11-0001, 11-0010, 11-0100, 11-1000) were inoculated from plates into liquid media containing 3.5 mM disodium succinate and 50 μg/mL kanamycin to grow cultures up to substantial biomass and ensure cell plasmid carriage. From here, cultures were diluted 1/64 in 5 mL liquid media containing 15 mM methanol and grown for 2 days and then subsequently diluted into new media. Additionally, we passaged genotype 11-0000 in liquid media with 3.5 mM succinate as the sole carbon source and lacking kanamycin as a control where we expected rapid loss of the plasmid. Every four days (2 transfers = 12 generations), subsamples were taken from the populations, and a dilution series was completed and plated on media with and without kanamycin to assess any change in frequency of plasmid-containing cells over time. A subsequent experiment involved 20 replicate populations of the 11-1010 genotype (strain CM3290, Fig. 6B).

## Results

### Many *mutation x pathway* epistatic interactions observed between alleles evolved with the *Foreign* pathway when present in *Native* and *Dual* pathway strains

Do the beneficial alleles that evolved in the context of the *Foreign* pathway also provide benefits with either the *Native* pathway alone or in the *Dual* pathway strains, or are there *mutation x pathway* interactions that change the sign of their effect? In this study, we sought to determine the fitness landscape that emerges upon joint consideration of beneficial mutations and gene gain and loss in the methylotrophic metabolism of *M. extorquens*. To do this, we constructed a series of strains that represent alleles with either just the *Foreign* or the *Native* pathway in isolation, as well as the *Dual* pathway strain (Table S1).

In contrast to the universally beneficial fitness effects observed for evolved alleles in the *Foreign* pathway context, introducing these alleles individually into the *Native* and *Dual* pathway backgrounds gave mixed results indicating many *mutation × pathway* epistatic interactions (Fig. 3, gray and orange edges; Fig. 4, faintly shaded and/or negative bars; Table S4). Only two alleles – *fghA*^evo^ or *gshA*^evo^ into *Dual* – were still individually beneficial. Alternatively, *GB*^evo^ was individually neutral and *pntAB*^evo^ was individually deleterious in either context. Fitnesses calculated from pairwise assays correlated well with exponential growth rates when strains grew individually, validating the fitness measure (Fig. S5A; strong positive correlation, R^2^=0.8928, p<0.0001).

**Figure 4.**
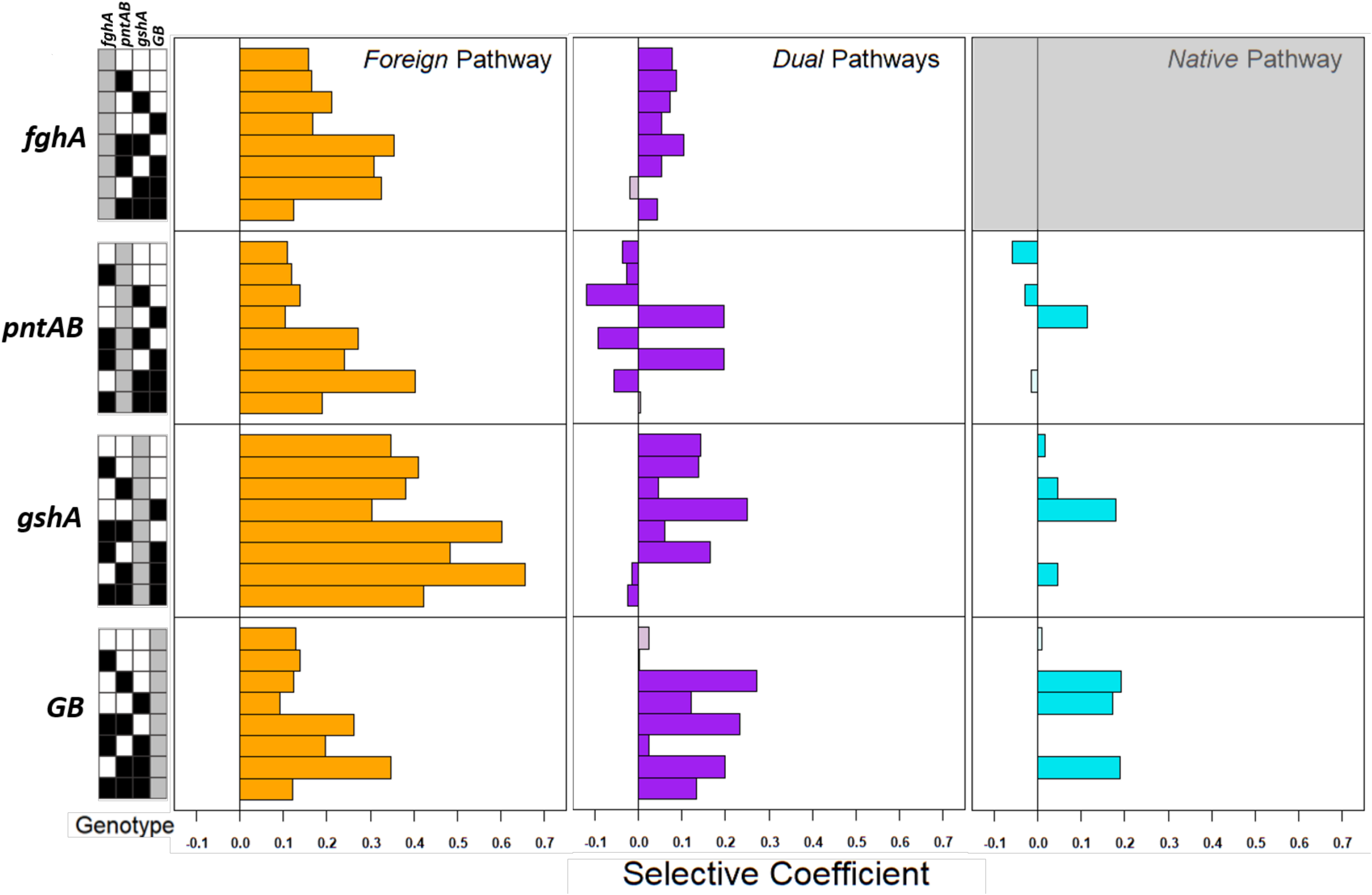
Comparison of the selective coefficients of mutational changes in different pathway backgrounds. For each panel, bars above the line are beneficial in effect, while those falling below are deleterious. The combinatorial genotype the focal allele is introduced into is depicted by the key below – ancestral alleles (‘0’) are coded in white, evolved alleles (‘1’) are coded in black, and the focal allele being introduced are color coded in an allele-specific manner (*fghA* = yellow, *pntAB* = blue, *gshA* = green, *GB* = red). The *Native* pathway *fghA* section is grayed out because these are not possible genotypes. The bars color intensity is weighted by the likelihood of draw, and those with intermediate draw probabilities (95% confidence intervals overlapping neutral) are lightly shaded.

### Abundant *mutation × mutation × pathway* interactions lead to *Native* and *Dual* fitness landscapes being incongruous with the *Foreign* pathway context

Moving beyond the individual effects of mutations in new pathways, does the manner in which they interact with each other in new metabolic settings – *mutation × mutation × pathway* interactions – further alter the fitness landscapes of combinations of these alleles in the *Native* and *Dual* contexts? The abundance of *mutation x mutation x pathway* interactions led us to examine whether these emerged due to idiosyncratic differences, or due to consistently different interactions of particular allele combinations. A few consistent allelic patterns emerged (Figs. 3 & 4). First, the *fghA*^evo^ allele remained beneficial in six of the seven additional *Dual* backgrounds that contained additional evolved allele(s). Second, all transitions to the *GB*^evo^ allele in the presence of *pntAB*^evo^ and/or *gshA*^evo^ were beneficial. This was surprising given that only one of these alleles in isolation provided strongly beneficial effects, and even then, in only one of the two pathway backgrounds (*gshA*^evo^ in the *Dual* background; Fig. 4). Third, *gshA*^evo^ was either beneficial or neutral when combined with other alleles, but the pattern of interaction differed between the *Dual* and *Native* contexts. Finally, the *pntAB*^evo^ allele exhibited abundant sign epistasis. It remained deleterious in half of the backgrounds containing the *gshA*^evo^ allele (the other four being neutral) yet was beneficial in half of the backgrounds with *gshA*^anc^ (and neutral for the other three).

### Epistatic interactions alter the fitness peaks and adaptive trajectories possible for combinations of alleles evolved with the *Foreign* pathway when tested in the *Native* or *Dual* contexts

Given the distinct individual and joint effects of alleles that were beneficial in the *Foreign* context when tested in the *Native* or *Dual* contexts, we next asked how this would alter adaptive outcomes that involved such mutations. To incorporate measurement uncertainty in our analysis, rather than just considering the mean effects, we simulated random draws from empirical fitness distributions for all tested combinatorial genotypes to see how likely it was that all mutational changes in a given trajectory were beneficial. In the *Foreign* landscape all trajectories were accessible (i.e., adaptive along the entirety of a series of transitions) as all mutations were universally beneficial (Fig. 3A, Table S6). In the *Native* and *Dual* pathway landscapes, however, this was not the case (Fig. 3 & S1, Table S6). It was much rarer that either the 10-x111 or 11-1111 genotypes were accessible as strictly accessible paths: only one of the six trajectories in the *Native* landscape and none of the 24 trajectories available on the *Dual* landscape were strictly accessible in even 50% of draws (Table S6). This indicates the existence of alternative local peaks in these landscapes. In the *Dual* pathway background, two genotypes (11-0011 and 11-1101) possess higher fitness values than 11-1111, and the overall fitness peak among this set is occupied by genotype 11-1101. This is also interesting because the genotypic combination producing a peak in the *Dual* landscape differs from the genotype producing a fitness peak on the *Native* landscape (10-x011), suggesting that different physiological conditions either favoring or disfavoring these combinations exist among them.

When we assess trajectories including transitions between only 3 or 4 genotypes (rather than 5 included in paths from 0 to 4 evolved alleles), we see that 12 are accessible with over 50% certainty, and 3 are accessible with 99% certainty (Fig. 3B, Table S7). In the *Foreign* landscape, all 24 of the possible trajectories to 01-1111 were universally beneficial with >99.9% certainty (Table S6). Conversely in the *Native* (5/6 paths accessible in < 50% of trials, 3/6 paths accessible in < 1% of trials) and *Dual* (24/24 paths accessible in < 50% of trials and 10/24 paths accessible in < 1% of trials) landscapes were universally less likely to be beneficial. Thus, while selection appeared to permit any adaptive trajectory to the peak of the *Foreign* landscape, limited paths and alternative peaks exist for these alternative backgrounds due to neutral and deleterious fitness effects.

### The cost of the plasmid carrying the *Foreign* pathway is impacted by epistatic interactions with evolved alleles present, and beneficial transitions from *Native* to *Dual* pathways are rare

Moving beyond considering the evolved alleles alone or in combination, we can also consider the loss of the plasmid carrying the degenerate, *Foreign* GSH pathway in *Dual* strains, and the resulting transitions to genotypes with only the *Native* pathway. The loss of pCM410 carrying the *Foreign* pathway was beneficial in the base (e.g., no evolved alleles) *Dual* pathway strain whether or not the *fghA*^evo^ allele was present (Fig. 5A). There are clear *mutation × mutation × pathway* interactions, however, as the magnitude of these costs varied with host genotype, (Fig. 5, Tables S5 & S8). The total carriage cost was generally lower if the plasmid contained the evolved *fghA*^evo^ allele that reduces expression, but there were substantial differences across allele backgrounds, with several showing more than a 50% reduction in costs due to the additional alleles. In one context (genotypes “1x-x010”), the combination of alleles present had a sufficiently strong effect to render the evolved version of the plasmid with *Foreign* pathway was beneficial. This general tendency for plasmid loss alters the accessible trajectories across both the *Dual* and *Native* landscapes, which can be visualized by the dominance of transitions that would transition *Dual* strains that contain both pathways to only containing the *Native* pathway (Fig. 5B).

**Figure 5.**
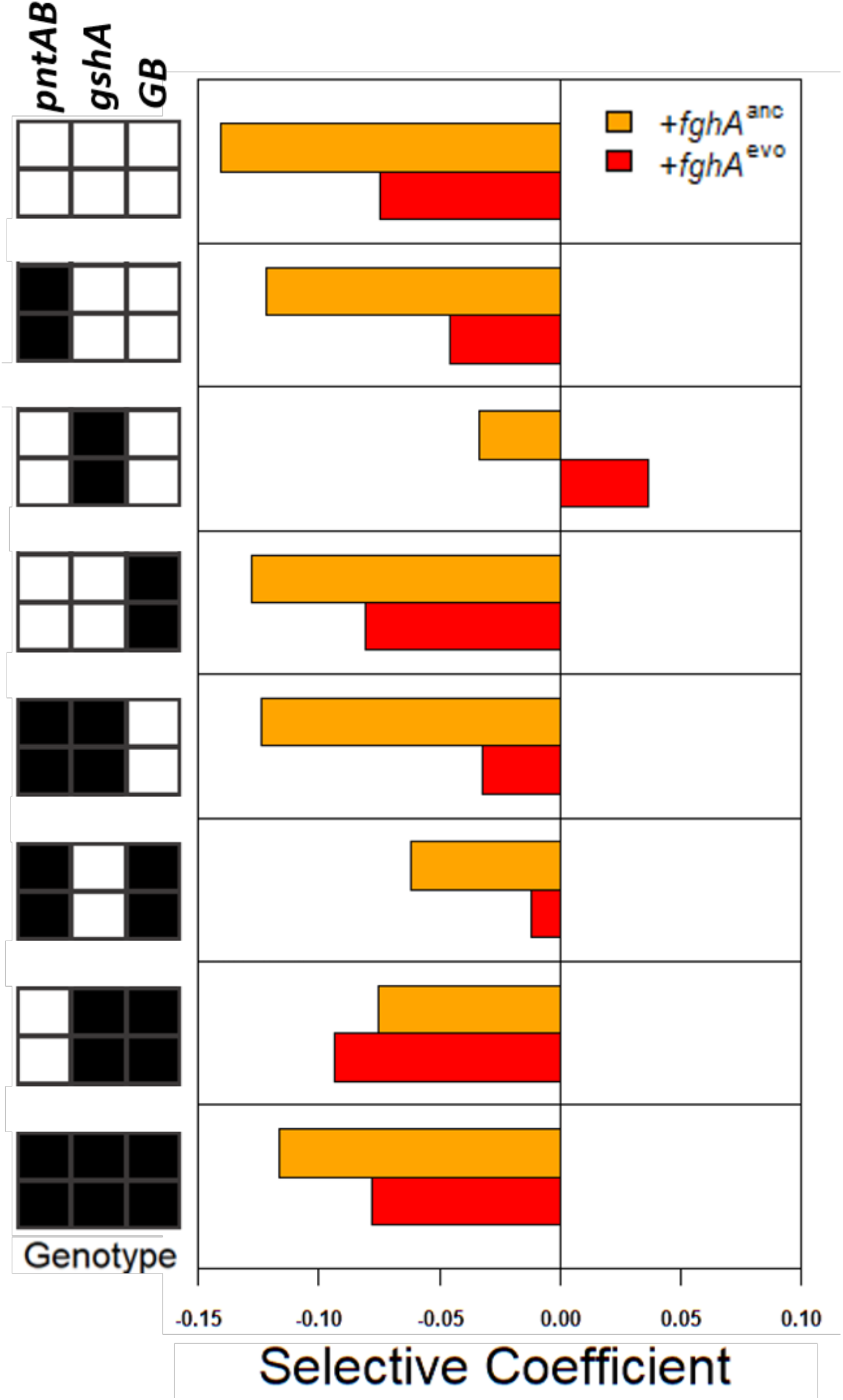
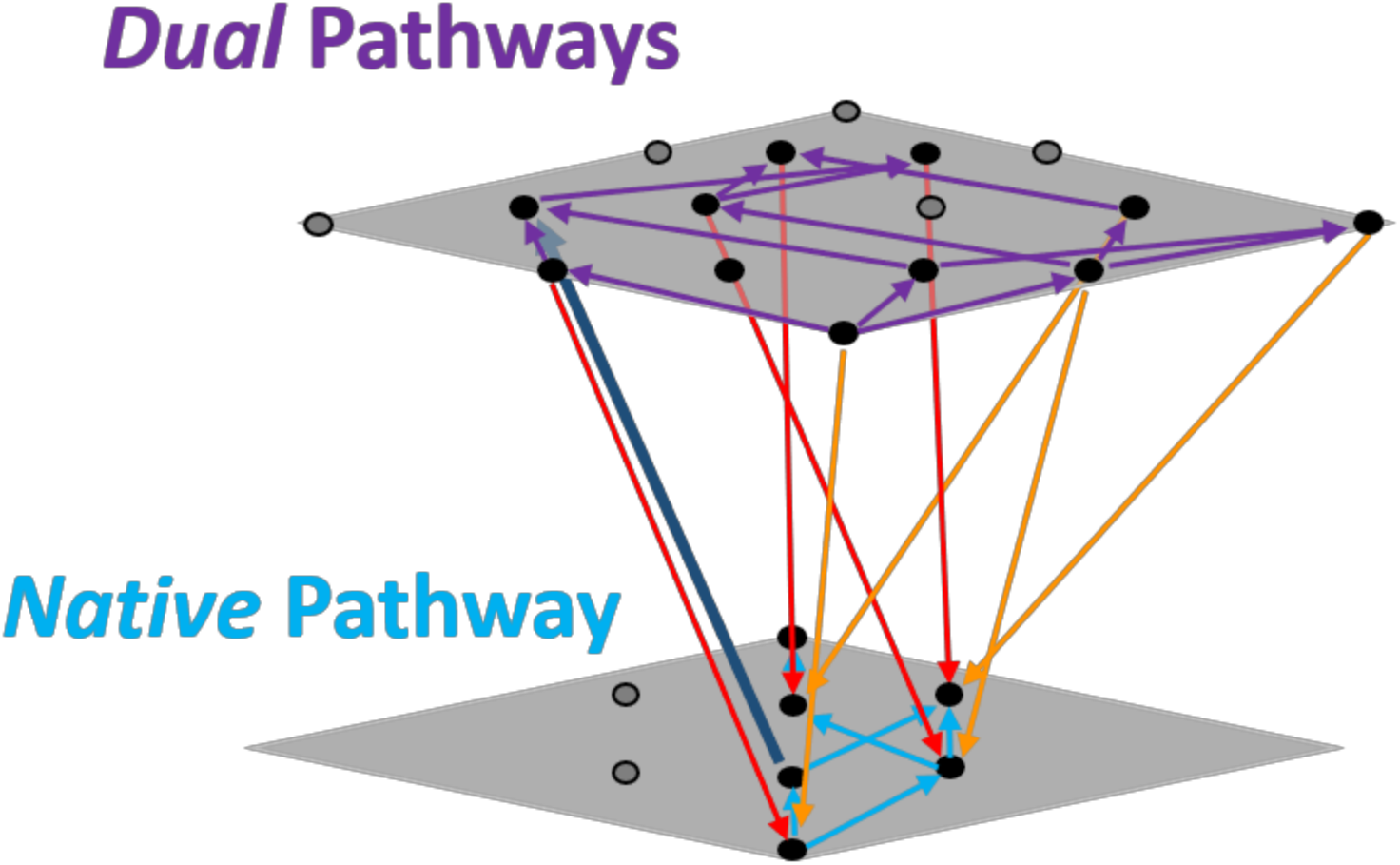
Factors impacting retention of a second degenerate metabolic pathway. **A)** Comparison of selective coefficients for inheriting the GSH pathway plasmid by *Native* pathway strains. The bars are color coded by which focal *fghA* allele is being introduced: *fghA*^anc^ is in orange and *fghA*^evo^ is in red. The specific genotypic background that the mutation is introduced into is indicated in the key at the bottom. Ancestral alleles (‘0’) are coded in white, evolved alleles (‘1’) are coded in black). **B)** Conceptual fitness landscape schematic depicting accessible paths within and between (by gene acquisition or loss) *Native* and *Dual* landscapes. Arrows indicate the direction of selection. Purple arrows indicate selectively favored transitions among the *Dual* landscape, and cyan arrows indicate selectively selectively favored transitions on the *Native* landscape. Orange and red arrows indicate selection favoring loss of degenerate pathway plasmid (bearing the *fghA*^anc^ and *fghA*^evo^ alleles, respectively), while the blue arrow indicates selection favoring plasmid retention.

### Rapid genotype-dependent loss of degenerate plasmid-borne glutathione-dependent pathway leads to rare retention and coevolution of multiple degenerate pathways

Do the plasmid costs inferred from competition experiments suggest that the degenerate pathway will simply be lost rapidly if acquired in this manner? Our competition-based results along with the inherent cost of pCM410 suggested that rapid loss would be expected in many *Dual* genotypic backgrounds if direct selection was removed (Figs. 3 & 5, Tables S5 & S8). To further address whether the second degenerate metabolic pathway would be incorporated into the metabolic network of recipient genotypes or lost if acquired via pCM410, we measured how well WT maintained the plasmid encoding the enzymes for the GSH pathway (genotype 11-0000’; Fig. 2). Over multiple passages in succinate-or methanol-based medium, we regularly monitored plasmid retention by quantifying viable cell counts on comparable solid medium with and without kanamycin (Fig. S7). Through this screen, we observed rapid and uniform loss of the plasmid from the basal *Dual* strain, independent of the carbon source included, contradicting our expectation that the pathway would be more expendable on a non-methylotrophic carbon source (succinate). This suggested that even when formaldehyde oxidation was required for growth (i.e., when growing on methanol), it was beneficial for WT to lose the degenerate pathway.

Despite the loss of the plasmid by the basal *Dual* strain, can any of the genotypic combinations examined enable the retention of a novel degenerate formaldehyde oxidation pathway? Because of the existence of non-deleterious mutational transitions in the *Native* and *Dual* backgrounds, we considered the possibility that additional evolved alleles or alternative environmental conditions might favor retention of a second degenerate formaldehyde oxidation pathway via epistatic interactions. To address this possibility, we tested additional *Dual* pathway genotypes for their maintenance or loss of the *Foreign* pathway plasmid via passage under methylotrophic growth conditions without antibiotic selection (Fig. 6). Several strains from the *Dual* pathway set of strains were tested to examine patterns of plasmid retention/loss as an indicator of how costly, neutral, or advantageous the presence of the plasmid-encoded *Foreign* pathway was, as well as the capacity for these strains to continue to coevolve with both pathways present. This included genotypes that do (e.g., 11-0100) and do not (e.g., 11-0010) possess negative fitness effects from carrying the second pathway (Figs. 5 & 6B). In fact, we saw that the relative rate of plasmid loss was consistent with the relative cost burdens that strains inherit after transitioning to *Dual* pathway genotypes (Figs. 5, 6A, & S7). The *GB*^evo^ strain, which had a similar plasmid maintenance cost to that of the base strain, saw a similar rate of plasmid loss to that strain. Furthermore, *Dual* strains with lower plasmid maintenance costs had concurrently reduced rates of plasmid loss (*gshA*^evo^ < *pntAB*^evo^ < *fghA*^evo^, Figs. 5 & 6A, Tables S5 & S8). This data provided further evidence that adding the second pathway did not provide net benefits beyond harboring the *Native* pathway alone and instead led to substantial fitness costs [54], further raising the question of how maintaining such degenerate paths could be favored by evolution (Figs. 5 & 6A).

**Figure 6.**
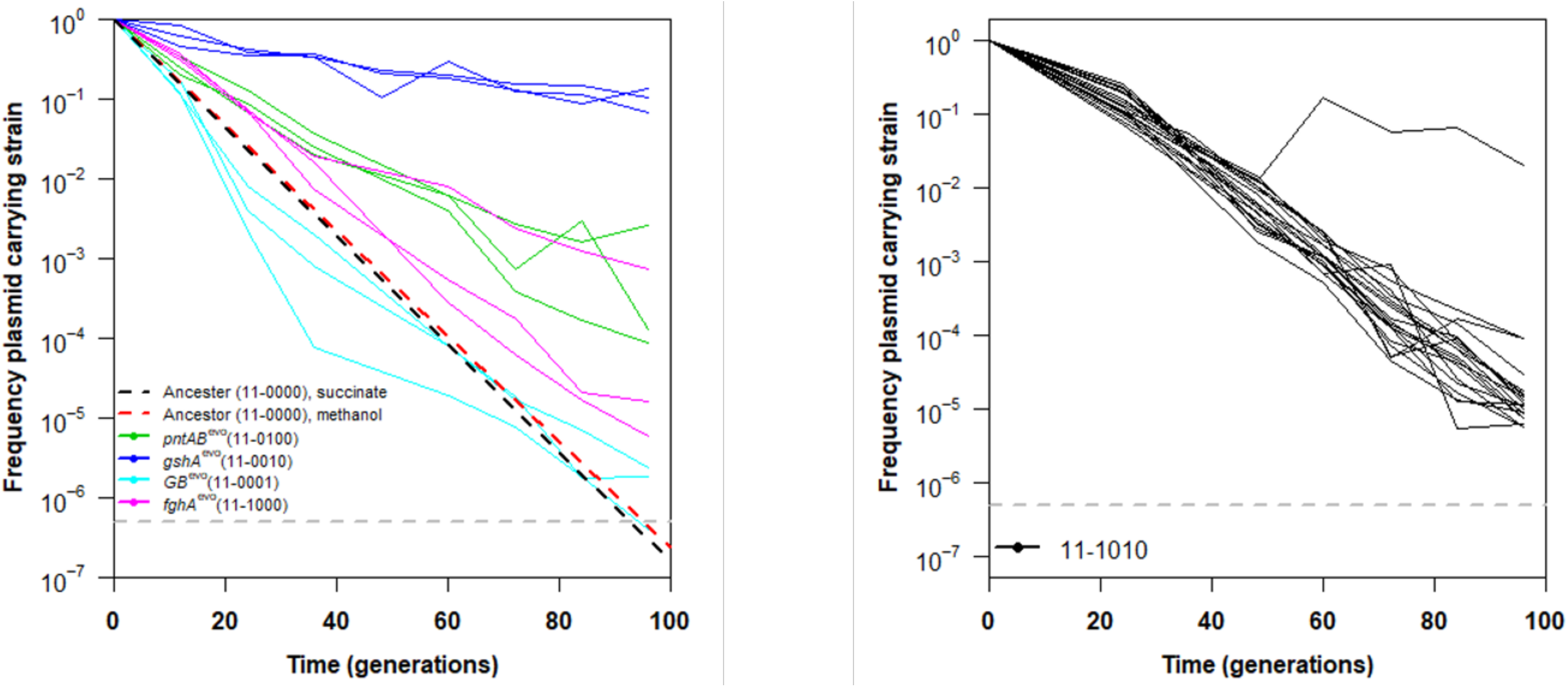
Plasmid carriage over time. Examination of plasmid pCM410 retention/loss over time was examined in strains of *M. extorquens* AM1 with evolved mutations identified in a previous study [19]. **A)** Each strain examined has a separate evolved mutation(s) in its background and was carried out in triplicate. Lines indicate the change of individual lineages and dots indicate the mean of the triplicate cultures for a given strain. Model fits (linear regression) for the rate of loss in the background strain CM3260 grown in both succinate (black) and methanol (red) media are also provided as dashed line for reference. **B)**. Plasmid carriage by genotype 11-1010. This strain was examined was carried out in biological replicate of N=20. Lines indicate the change of each individual lineage and dots indicate the mean of the replicate cultures. Both experiments were carried out in Hypho media with relaxed selection for the plasmid (no antibiotic present). The dashed gray line in each panel indicates the experimental limit of detection for the assay.

To give adaptation of degeneracy the greatest opportunity possible with these strains, we propagated 20 populations of the 11-1010 genotype, which was the one example where the presence of the plasmid with the *Foreign* pathway was beneficial. Most of these populations also experienced rapid loss of the plasmid, however, we saw reversal of plasmid loss around 50 generations into the experiment for one population (Fig. 6B). At this point, the frequency of the plasmid carrying strain increased by an order of magnitude then decreasing at a pace much slower than all other lineages. The reduced rate of plasmid loss in *gshA*^evo^-containing lineages suggests that they have the capacity to maintain the degenerate pathway long enough for adaptive mutations to arise that maintain the associated degeneracy/novel functions [63].

## Discussion

This study sought to address an open question: how might the evolution of a novel degenerate pathway for a given metabolic process (such as encountered in nature through HGT) be impacted in different genetic (and resulting physiological) contexts? In particular, what are the challenges associated with evolving to acquire and retain multiple, degenerate pathways that provide analogous functions for the cell? Combining the *Native* dH_4_MPT and *Foreign* GSH pathways for formaldehyde oxidation might seem like a non-starter: introduction of the *Foreign* pathway to generate the *Dual* pathway strain was found to be deleterious [19,20]. Indeed, propagation of the original WT strain with *Foreign* pathway led to rapid loss (Fig. S7). A strain with the *Foreign* pathway alone rapidly evolved improved growth, but previous work left unaddressed whether this evolution would improve or further limit opportunities for coexistence of the two pathways. Such adaptive changes might have ameliorated the costs associated with maintaining the *Foreign* pathway or, via tradeoffs, led to antagonism with the original, *Native* pathway, such that each improvement in the context of the *Foreign* pathway alone would come at a cost when coupled with the *Native* pathway. Critical to understanding these adaptive possibilities is to map out a broader picture of epistatic interactions, moving from evolved alleles (*mutation × mutation*), or alleles in a metabolic context (*mutation × pathway*), to consider the combined *mutation × mutation × pathway* interaction network. These interactions produce a broader view of the fitness landscape that determines the gain and loss of pathways (Fig. 1).

Whereas the fitness effects of the evolved alleles tested were universally beneficial across genotypes in the original *Foreign* context, we observed rampant *mutation × pathway* and *mutation × mutation × pathway* effects when these were placed in the *Dual* and *Native* contexts. None of the evolved alleles had universal selective effects across genotypes tested. The simplest set of hypotheses would have been that alleles would be universally beneficial in the *Dual* and *Native* contexts if they improve growth on methanol (or other environmental factors in the selective regime) in a manner independent of the pathway(s) present, they would be universally neutral if they aided the *Foreign* strains in using the GSH pathway in a manner irrelevant to functioning of the *Native* pathway, or universally deleterious if they compromised the function of the *Native* pathway in order to improve use of the *Foreign* pathway. Instead, the *mutation × mutation × pathway* interactions were prevalent and substantial. Each of the three alleles had their qualitative effect (i.e., beneficial vs. neutral vs. deleterious) altered by the presence of other alleles, and these patterns tended to be inconsistent between the *Dual* and *Native* pathway contexts.

The wide range of epistatic effects observed in the *Dual* and *Native* contexts often ran counter to expectations based upon the known physiological connections between them. As described, the *Foreign* GSH pathway depends upon a distinct C_1_ carrier, GSH rather than dH_4_MPT, and requires involvement of transhydrogenase (encoded by *pntAB*) to generate NADPH and recycle NADH to NAD^+^ for further use by the GSH pathway. Correspondingly, it has been observed that the ratio of NADH and NADPH is disturbed in the *Foreign* strain compared to the *Native* WT [22]. Furthermore, GSH is active as a C_1_ carrier when in the reduced thiol form, a process catalyzed by glutathione reductase, which requires NADPH as its source of reducing equivalents. We therefore hypothesized a positive interaction between *gshA*^evo^ and *pntAB*^evo^, which respectively upregulate expression of the genes contributing to GSH and NADH production, at least in the *Dual* pathway strain. Instead, the fitness cost of *pntAB*^evo^ was greater when introduced into backgrounds already containing *gshA*^evo^, and vice versa (P=0.005, unpaired t-test). In light of this observation, it appears that increased transhydrogenase activity might be expected to be universally deleterious in the *Native* pathway context due to the dH_4_MPT pathway already being capable of generating either cofactor, such that transhydrogenase activity could lead to a futile cycling due to coupling to dissipation of the transmembrane proton gradient. In contrast, whereas *pntAB*^evo^ was deleterious as the sole additional allele introduced, in many of the other backgrounds it was neutral, and, in one case, even beneficial. Finally, the *GB*^evo^ allele is a composite of six mutations, one of which (the large deletion) has been directly demonstrated to be beneficial in the *Native* pathway context of the WT strain during growth on methanol, or even other growth substrates [21]. Here we observed that *GB*^evo^ behaved as a neutral allele when introduced alone into *Dual* or *Native* contexts, but in many of the combinations could be strongly beneficial, indicating synergistic epistasis with other evolved alleles (particularly *pntAB*^evo^ or *gshA*^evo^). At this point, the physiological basis of these many examples of sign epistasis within the *mutation x mutation x pathway* interactions remains unclear, but leads to caution in assuming that either the individual effects of mutations in new pathway contexts will hold when alleles are combined, or that the known *mutation x mutation* interactions observed in one pathway context will hold when present in a new metabolic setting.

Given the extent of epistasis observed, would we expect evolution to favor similar beneficial mutations across the contexts of the *Native*, *Foreign*, and *Dual* pathway genotypes? If only individual mutational effects are considered, there was little congruence between these landscapes. But due to *mutation x mutation × pathway* epistatic interactions, the fitness peaks on each of the *Dual* and *Native* landscapes had many overlapping alleles with the optimal alleles in the *Foreign* context. The frequent neutral or deleterious effects observed for evolved alleles in the *Dual* and *Native* landscapes, however, would generate a much more restricted set of paths to these peaks. Interestingly, the historical sub-trajectory known for the *Foreign* pathway population studied (*gshA*^evo^ then *fghA*^evo^ then *pntAB*^evo^) [23] would not be selectively accessible in either the *Dual* or *Native* pathway contexts.

The overall tendency for the *Foreign* pathway to be deleterious in the *Dual* pathway context suggests that in many cases degeneracy would simply resolve to having just the *Native* pathway. This grim expectation for the *Foreign* pathway should be tempered for several reasons. First, the magnitude of the cost of the *Foreign* pathway was quite flexible, with nearly all allele combinations exhibiting a lower cost than the ancestral plasmid in the ancestral *Dual* strain. It was unsurprising that the *fghA*^evo^ allele, which has been shown to reduce protein expression costs [19,20,54], generally reduced the cost of the degenerate pathway considerably. Beyond this, however, the precise combinations of alleles in the *Dual* pathway context mattered greatly. In one case, the evolved plasmid (e.g., *fghA*^evo^) in a genome with *gshA*^evo^ was even found to impart a benefit (Fig. 5). Direct tests of plasmid maintenance confirmed that rates of loss for this pathway were strongly dependent on pathway costs, which vary as a direct result of epistatic interactions (Figs. 5A & 6).

One population showed that evolution might act to maintain both pathways (Figs. 5B & 6B, [64]). From a total of 20 initial replicates, one population exhibited an increase in plasmid carriage, followed by a slow decline. Whereas both the rate of plasmid loss across cells (e.g., lack of segregation) and selection against plasmid containing cells [54,65] will contribute to the exponential decline in their frequency in a population, a rise in frequency requires either plasmid exchange, which is absent in this system, or a selective benefit large enough for that lineage to have a fitness higher than the average fitness in the population at that time. Given the known prevalence of clonal interference in populations with the *Foreign* pathway [66], at least when it is alone in the cell, these dynamics are consistent with that lineage emerging to compete with other high fitness lineages also segregating in the population. Further work will be required to isolate this lineage and determine the genetic basis of the first steps towards adapting towards degeneracy in that context. Although there was this one hopeful example, the majority of populations from this ancestor did ultimately lose the plasmid encoding the GSH pathway (Fig. 5B).

While our results generally illustrate the many challenges with adapting to utilize a new, *Foreign* metabolic pathway when present together with the *Native* pathway in a *Dual* context, there are factors that might increase the potential for redundancy to emerge. First, many broad-host-range plasmids encode functions to aid their carriage despite imposed fitness costs [67–71]. Second, *Foreign* pathways introduced directly into the chromosome would enjoy several advantages: stable segregation, generally smaller costs (if any), and lower copy number with would reduce gene dosage. Consistent with this, integration events of the pCM410 plasmid with host genome were incredibly common during adaptation of the *Foreign* pathway populations [20,66]. Third, pathways with low or appropriately-regulated expression may be expected to have lower initial costs; the plasmid used here and in the previous studies with this system had high, relatively constant level of expression [19,20,54,72]. This study adds to this list a fourth possibility, epistatic interactions between beneficial mutations and the pathway(s) present that can reduce or eliminate maintenance costs. Some of these may be compensatory mutations that alleviate plasmid maintenance costs [73,74], but others may be direct mediation of the physiological effects of a *Foreign* pathway. Given the abundant evidence of the importance of HGT as an avenue for evolution in microbes [75–79], clearly these hurdles have been overcome in a variety of cases. Very often this has led to strains with tremendous degeneracy, containing several unrelated versions of functional modules that provide similar benefits to the cell [11,80,81]. This natural variety, in the light of the results described here, should motivate future evolution experiments that focus on the gain of redundant pathways; like gene duplication, conflicts and redundancies may resolve in diverse ways with large potential consequences for evolutionary innovations.

## Acknowledgments

We thank and acknowledge funding from NSF to C.J.M. (MCB-1714949) and J.A.D. (MCB-1714550), from NIH to C.J.M. (R01 GM078209), from the NSF-funded BEACON Center for the Study of Evolution in Action (parent award DBI-0939454) to E.L.B. and C.J.M, from the University of Idaho Office of Undergraduate Research to C.J.R., and NSF REU support for N.V.E (DBI-1757826). We thank J. Bazurto (https://orcid.org/0000-0001-9012-2260) and members of the Marx lab for critical reading and commentary on the manuscript.

## Tables

**Table S1**. Strains used in this study. In order to simplify description of the complex genotypes in the text, pertinent genotypic aspects of the strains are listed as bitstring genotypes, as previously ([19], also see Fig. S1). To describe pathway background, “0” and “1” are utilized for the absence or presence of both the dH_4_MPT and GSH pathways (in that order). For the additional evolved alleles, a “0” represents the ancestral allele (“^anc^”, i.e. present at the start of the evolution experiment in [19]) and a “1” represents the evolved form of the allele (“^evo^”) that emerged in the sequenced isolate CM1145 from population F4. The ordering of these alleles is as such: *fghA* (encoding formyl-GSH hydrolase and hydroxymethyl-GSH dehydrogenase)/*pntAB* (transhydrogenase)/*gshA* (γ-glutamylcysteine synthetase)/*GB* (“Genetic Background”, a composite of six additional evolved mutations). For *Native* pathway strains, an “x” is included for the *fghA* allelic state because it is absent. As an example, “10-x100” represents a strain with a genotype in which the *Native* pathway is present and the *Foreign* pathway is absent and contains an evolved *pntAB* allele and ancestral *gshA* and *GB* alleles.

**Table S2**. Relative fitness values for strains assessed in competition assays.

**Table S3**. Probability of beneficial fitness draws and selective coefficients for *Foreign* pathway strains.

**Table S4**. Probability of beneficial fitness draws and selective coefficients for *Native* and *Dual* pathway strains.

**Table S5**. Probability of beneficial fitness draws and selective coefficients for transitions from *Foreign* or

*Native* to *Dual* pathway strains.

**Table S6**. Probability of accessible full trajectories from fitness draws.

**Table S7**. Probability of accessible partial trajectories from fitness draws for *Dual* pathway strains.

**Table S8**. Side by side fitness (vs. WT) comparisons for strains with methanopterin pathway present.

---

**Figure S1**. Primer on **A)** genotypes used in this study, **B)** transitions on fitness landscapes, and **C)**in/accessible evolutionary trajectories. A) Genotypes can be referenced/visualized both in terms of box diagrams (top) or as a series of bitstring numerals (bottom). From left to right, genotype boxes represent 1 & 2) pathway background, first dH_4_MPT and then GSH, 3) *fghA*, 4) *pntAB*, 5) *gshA*, and 6) *GB*. Coloration of the boxes depicts their status. Pathway backgrounds are filled with black when present and left unfilled when absent. The four other boxes may be colored in white (ancestral allele state) or black (evolved allele state). Bitstring sequences may contain “0” (for ancestral allele), “1” (for evolved allele), or “x” in cases where a particular allele is absent (specific to *fghA* in the *Native* pathway background). B) A focal genotype is separated by single mutations from its nearest mutational neighbors. The mutational transitions may be beneficial (green), deleterious (orange), or neutral (within +/-1% *S*, depicted in gray), the effect gauged by the probability of each class of effect occurring over many simulated draws. The weighting of the arrows reflects those probabilities, with thicker arrows more likely to be the type of effect it is color-coded by. C) Accessible and inaccessible evolutionary trajectories. Each fitness landscape (top) is composed of many trajectories within it. Those that are accessible (a subset of these shown on lower left) are a connected series of beneficial mutational transitions to an endpoint genotype. Inaccessible trajectories (a subset shown on the lower right) are those paths that are blocked because they involve steps that are neutral (gray) or deleterious (orange) that make it unfavored to be selected for. Where probabilities on given trajectories are measured, fitness values for each genotype along that trajectory are simultaneously drawn and that instance is considered accessible only if all mutational transitions along it are beneficial.

**Figure S2**. Fitness of *Foreign* pathway strains. **A)** Comparison of fitness results for *Foreign* pathway strains in those measured in this study to those from Chou *et al*. 2011 [19]. The inset graph includes all genotypes bearing strong correlations to fitness values reported in [19]. **B)** Relative fitness values for all *Foreign* pathway strains used in this study.

**Figure S3**. Comparison of fitness estimates derived from competition experiments utilizing the ancestor (01-0000) or wild-type strains (10-x000) for strains containing either the *Native* or *Dual* pathways. Because these strains were much fitter than the *Foreign* pathway strains, the corresponding fitness values from those competitions were much than those from WT competitions. However, the overall correspondence between these values was consistent (R^2^ = 0.68). The line of best fit is displayed.

**Figure S4.**Probability of a beneficial effects (> 0% represented by circles and >1% represented by triangles) in simulated draws from fitness distributions for examined mutational changes as a function of the mean selective coefficient for that change.

**Figure S5**. Growth analysis of strains from *Native* pathway set. Cell cultures of all genotypes from the *Native* pathway set, totaling 8 strains, were grown in triplicate in Hypho media with methanol as the sole carbon source. **A)** displays the relationship between the growth rates obtained from the growth experiments in **A)** and the relative fitness values for each strain displayed in **Figure 3**. Error bars represent standard error of the mean for 3 biological replicates. **B)** Monitored growth by absorbance at 600 nm (A600) over time. The line displayed is the result of a linear regression these values, with an R^2^=0.8928 (p=0.000123).

**Figure S6**. Histograms of selective coefficients calculated from fitness draws for mutational transitions. Distributions or all within-landscape transitions (*Dual* → *Dual*, for example) are displayed, along with mean value (also represented by a black vertical line) and the probability that a given draw will be beneficial, deleterious, or neutral (within 1% *S*). The title for each panel corresponds to the specific mutational transition, with both connected genotypes listed in bitstring notation. When visible on the displayed range, a completely neutral value (0%) is marked by a vertical red line and dashed red lines indicate +/−1% *S*.

**Figure S7**. Plasmid carriage over time. Examination of plasmid pCM410 retention/loss over time was examined when introduced into wild-type *M. extorquens* AM1(resulting in strain CM3260, genotype ’11-0000’). The experiment was carried out in Hypho media with relaxed selection for the plasmid (no antibiotic present). Regardless of the specific carbon source used, we observed rapid and constant loss of the plasmid over time, suggesting strong fitness costs for its carriage by this genotype.

**Figure S8.** *Foreign* fitness landscape depicting fitness values against the Ancestor strain. The same landscape displayed in Figure 3 for the *Foreign* pathway strains is displayed here but with fitness values reflective of their competitive outcomes against the *Foreign* ancestor control strain, CM1232.

