## Supplementary Figures for "Epistatic interactions shape the interplay between beneficial alleles and gain or loss of pathways in the evolution of novel metabolism"

### Figure S1A. Genotypic notation

Box genotype representations:

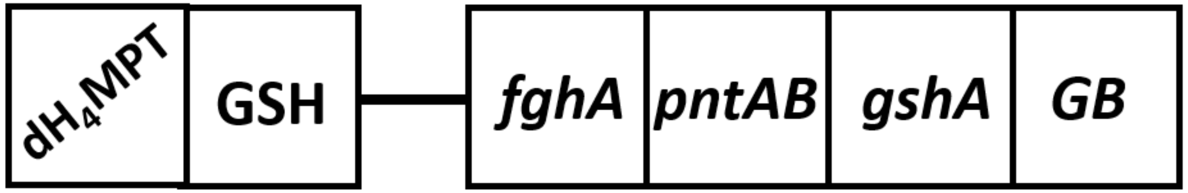


Bitstring genotype representations:

| | $dH_4MPT$ pathway | GSH Pathway | *fghA* | *pntAB* | *gshA* | *GB* |
|---|---|---|---|---|---|---|
| Ancestral ("anc"): | NA | NA | 0 | 0 | 0 | 0 |
| Evolved ("evo"): | NA | NA | 1 | 1 | 1 | 1 |
| Absent: | 0 | 0 | x | NA | NA | NA |
| Present: | 1 | 1 | NA | NA | NA | NA |

#### Examples: (bitstrings in parentheses)

Ancestor: (01-0000)

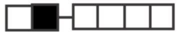

Wild-type: (10-x000)

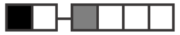

*Dual* Ancestor: (11-0000)

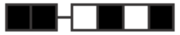

*Foreign GB*$^{evo}$: (01-0001)

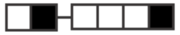

*Native pntAB*$^{evo}$ *gshA*$^{evo}$ : (10-x110)

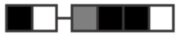

*Dual fghA*$^{evo}$ *pntAB*$^{evo}$ *gshA*$^{evo}$ *GB*$^{evo}$ : (11-1111)

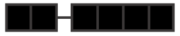

### Figure S1B. Fitness Landscape Transitions

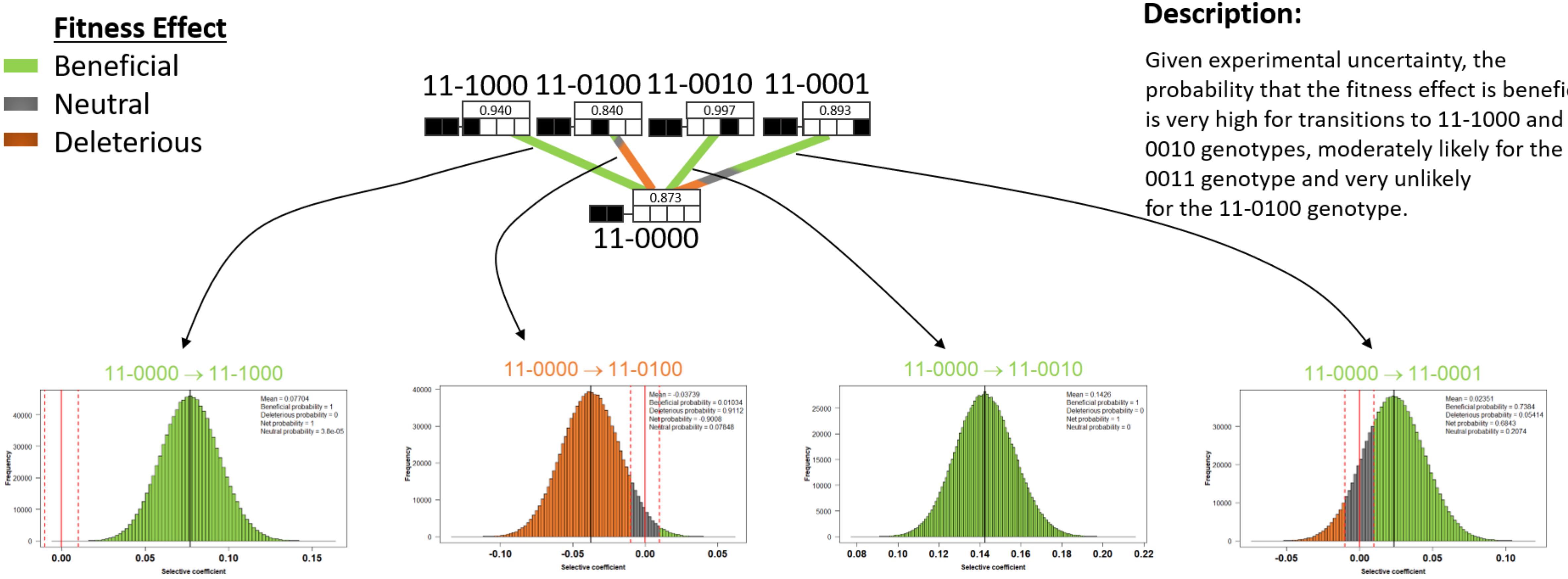


#### Description:

Given experimental uncertainty, the probability that the fitness effect is beneficial is very high for transitions to 11-1000 and 11-0010 genotypes, moderately likely for the 11-0011 genotype and very unlikely for the 11-0100 genotype.

### Figure S2.

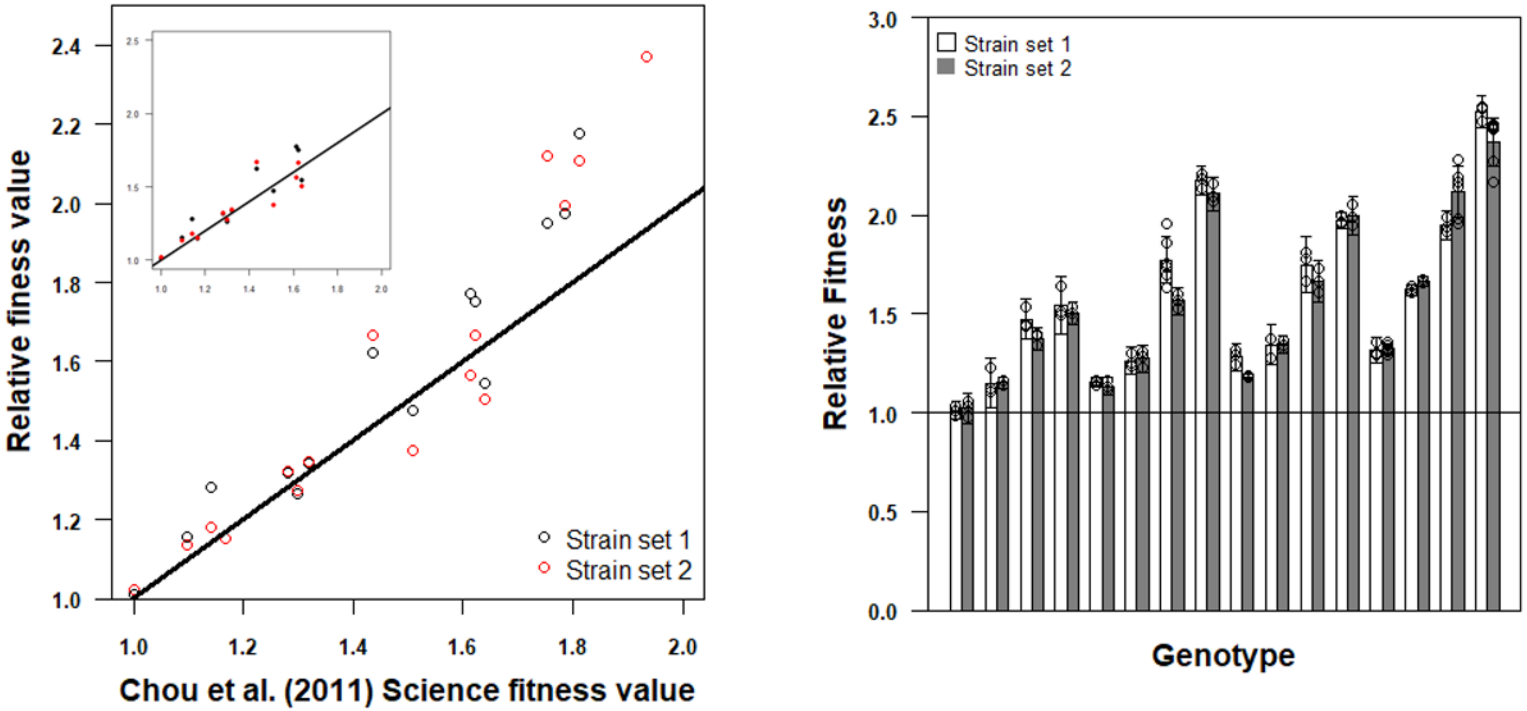
Relative finess value
Chou et al. (2011) Science fitness value
Strain set 1
Strain set 2
Relative Fitness
Genotype
Strain set 1
Strain set 2

### Figure S3.

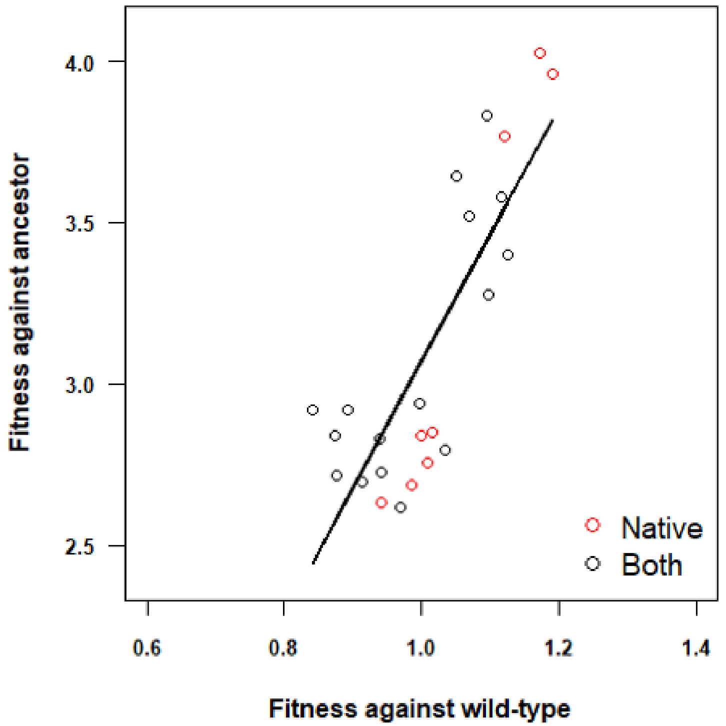

Fitness against ancestor
4.0
3.5
3.0
2.5
0.6
0.8
1.0
1.2
1.4
Fitness against wild-type
Native
Both

### Figure S4.

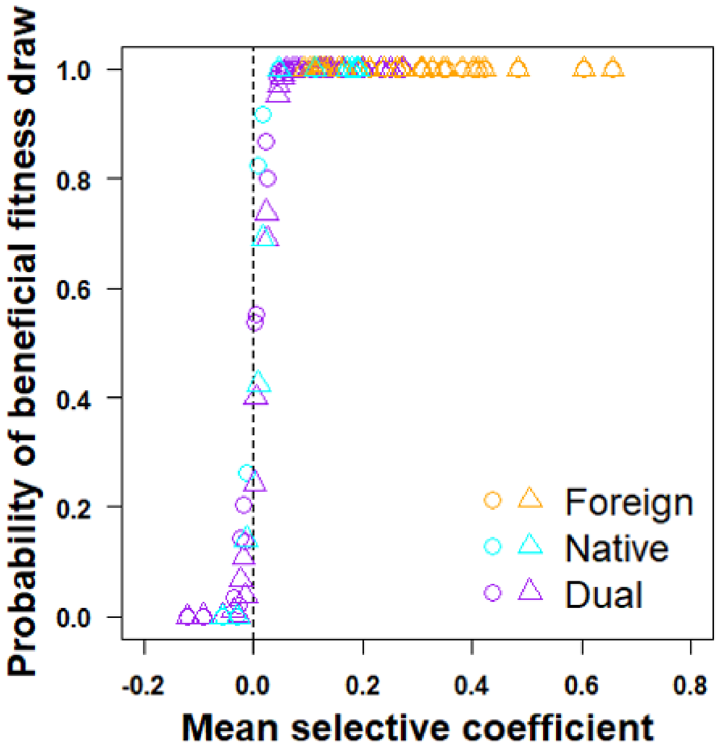

Probability of beneficial fitness draw
1.0
0.8
0.6
0.4
0.2
0.0
-0.2
0.0
0.2
0.4
0.6
0.8
Mean selective coefficient
Foreign
Native
Dual

### Figure S5.

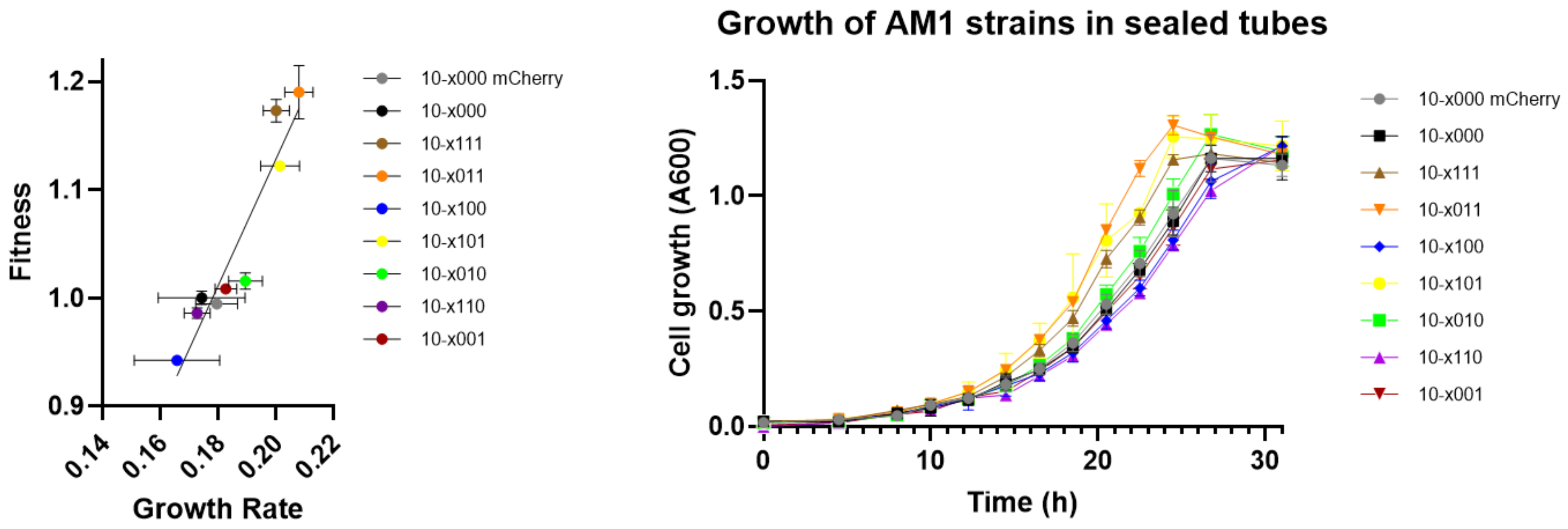
Fitness
Growth Rate
1.2
1.1
1.0
0.9
0.14
0.16
0.18
0.20
0.22
10-x000 mCherry
10-x000
10-x111
10-x011
10-x100
10-x101
10-x010
10-x110
10-x001
Growth of AM1 strains in sealed tubes
Cell growth (A600)
Time (h)
1.5
1.0
0.5
0.0
0
10
20
30
10-x000 mCherry
10-x000
10-x111
10-x011
10-x100
10-x101
10-x010
10-x110
10-x001

**Figure S6.**
***Foreign***

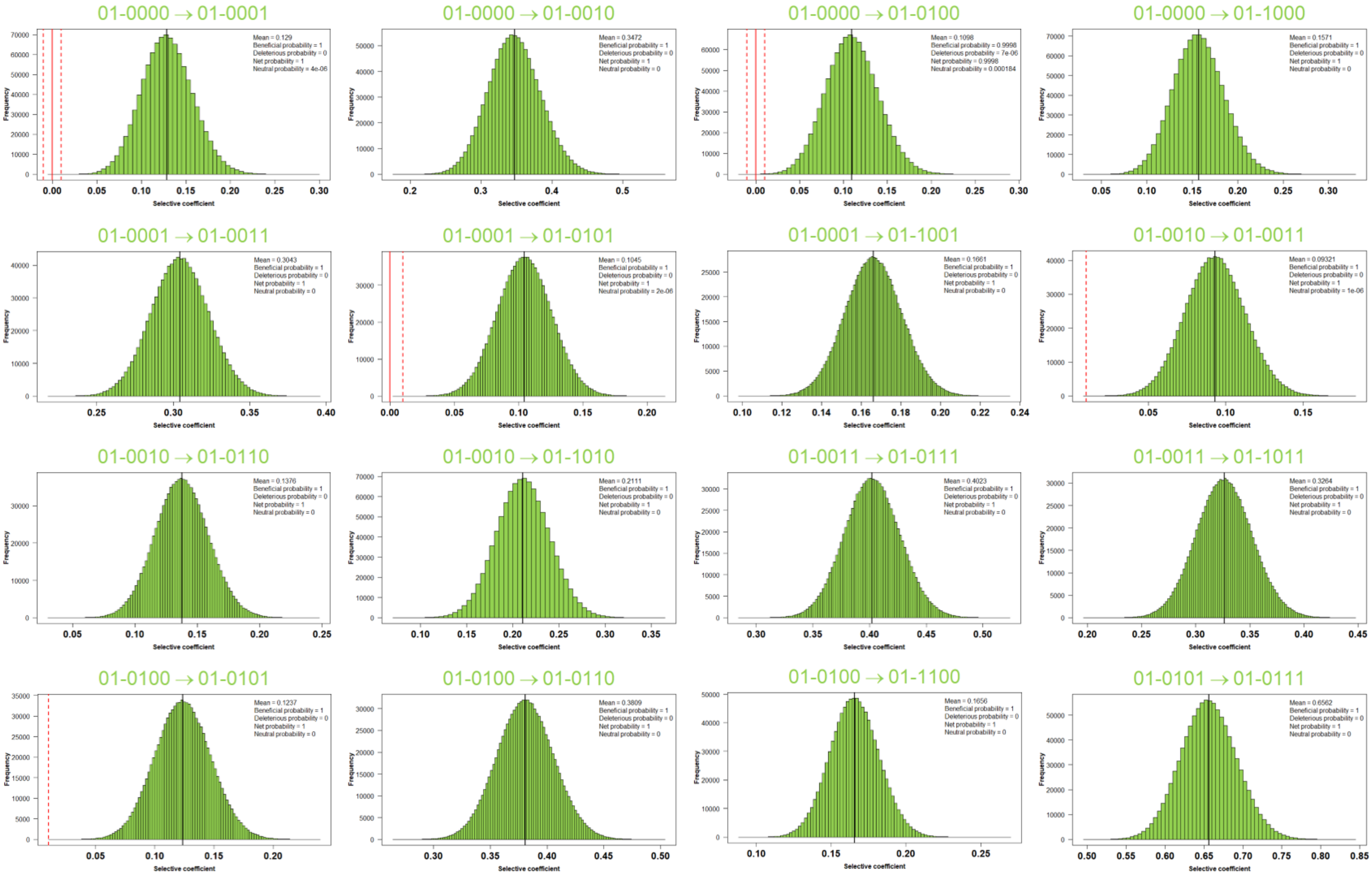

**Figure S6.**
***Foreign***
**(con't)**

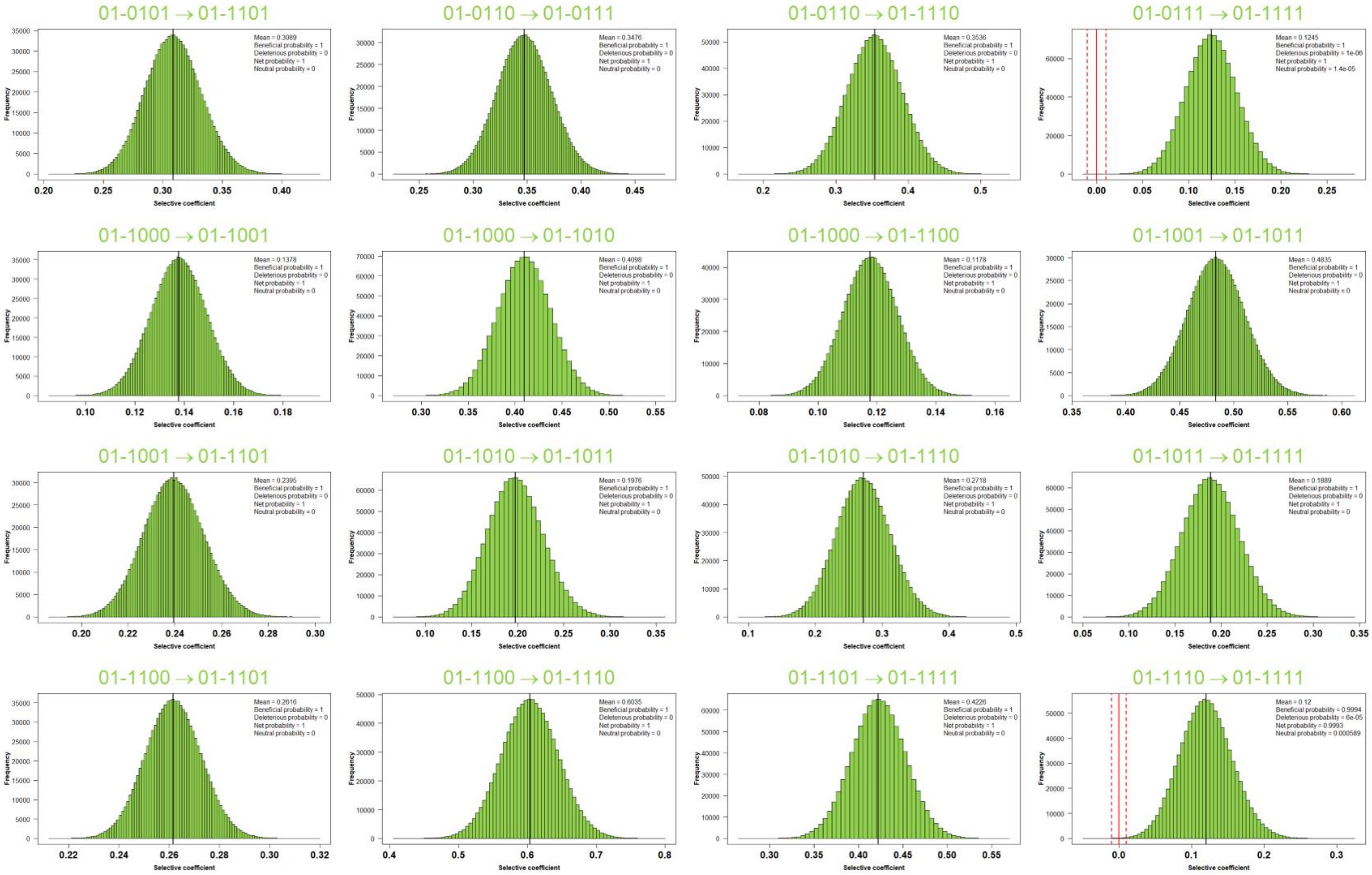

**Figure S6.**
***Dual***

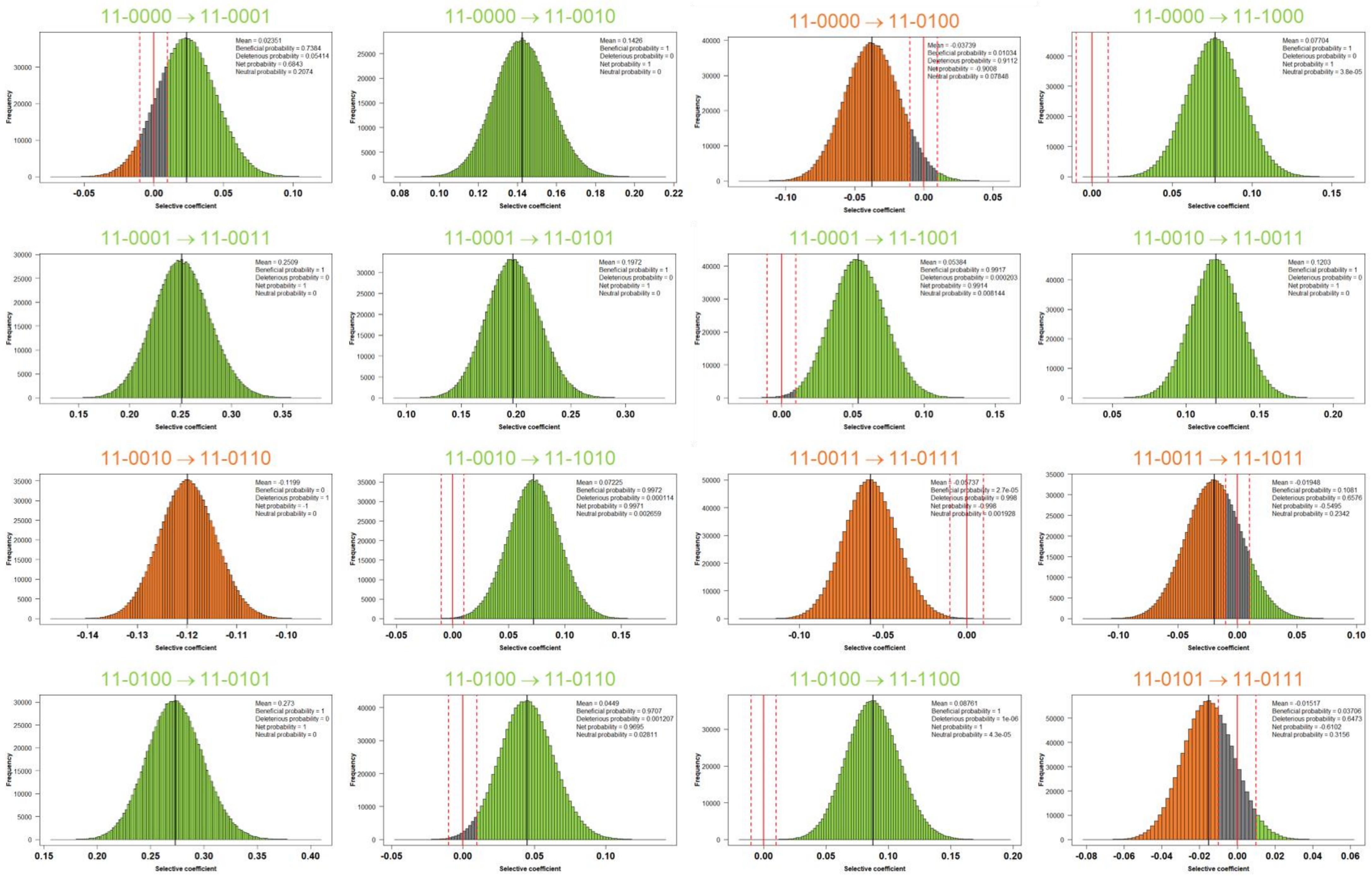

**Figure S6.**
***Dual***
**(con't)**

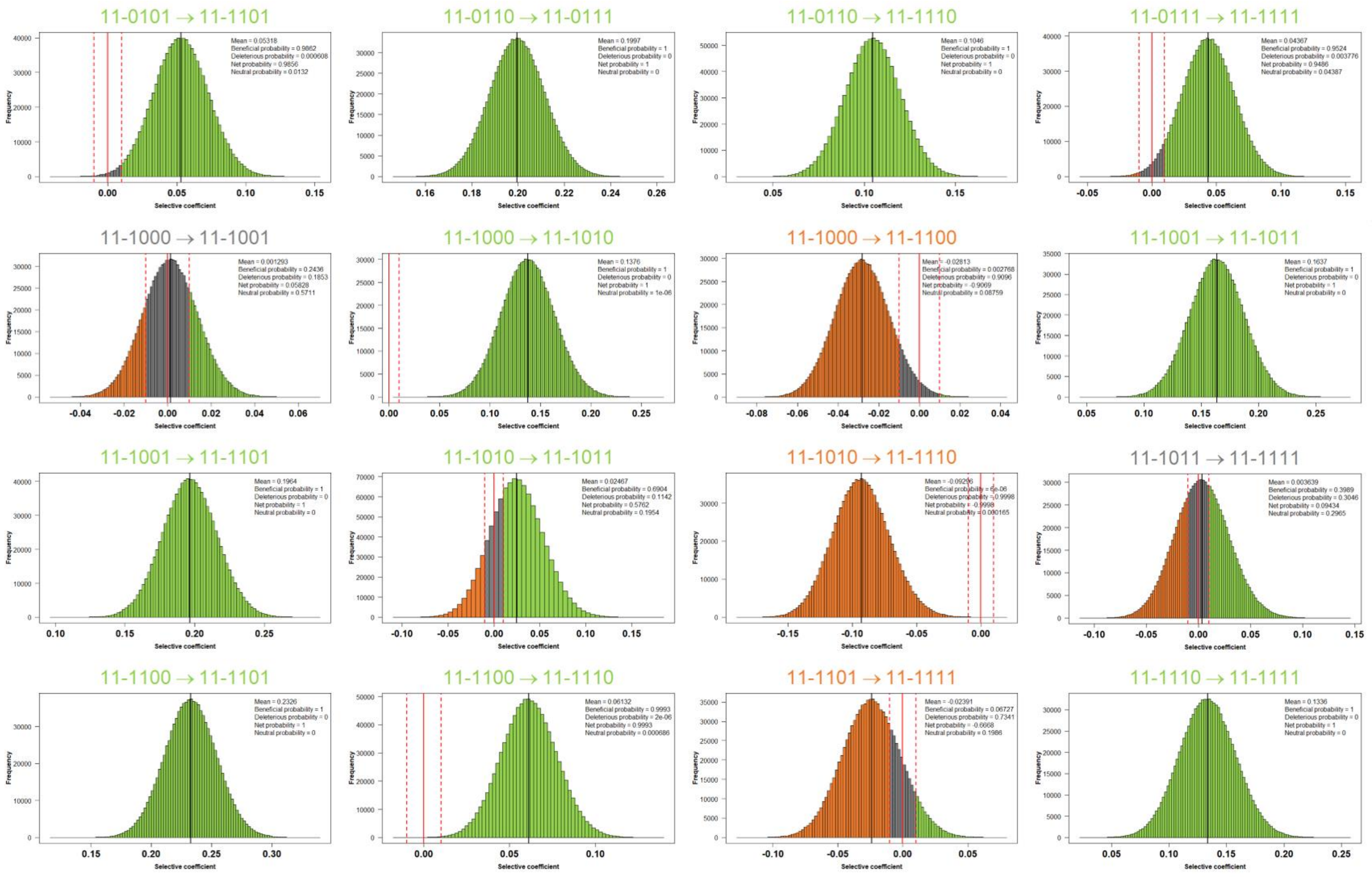

### Figure S6.
#### *Native*

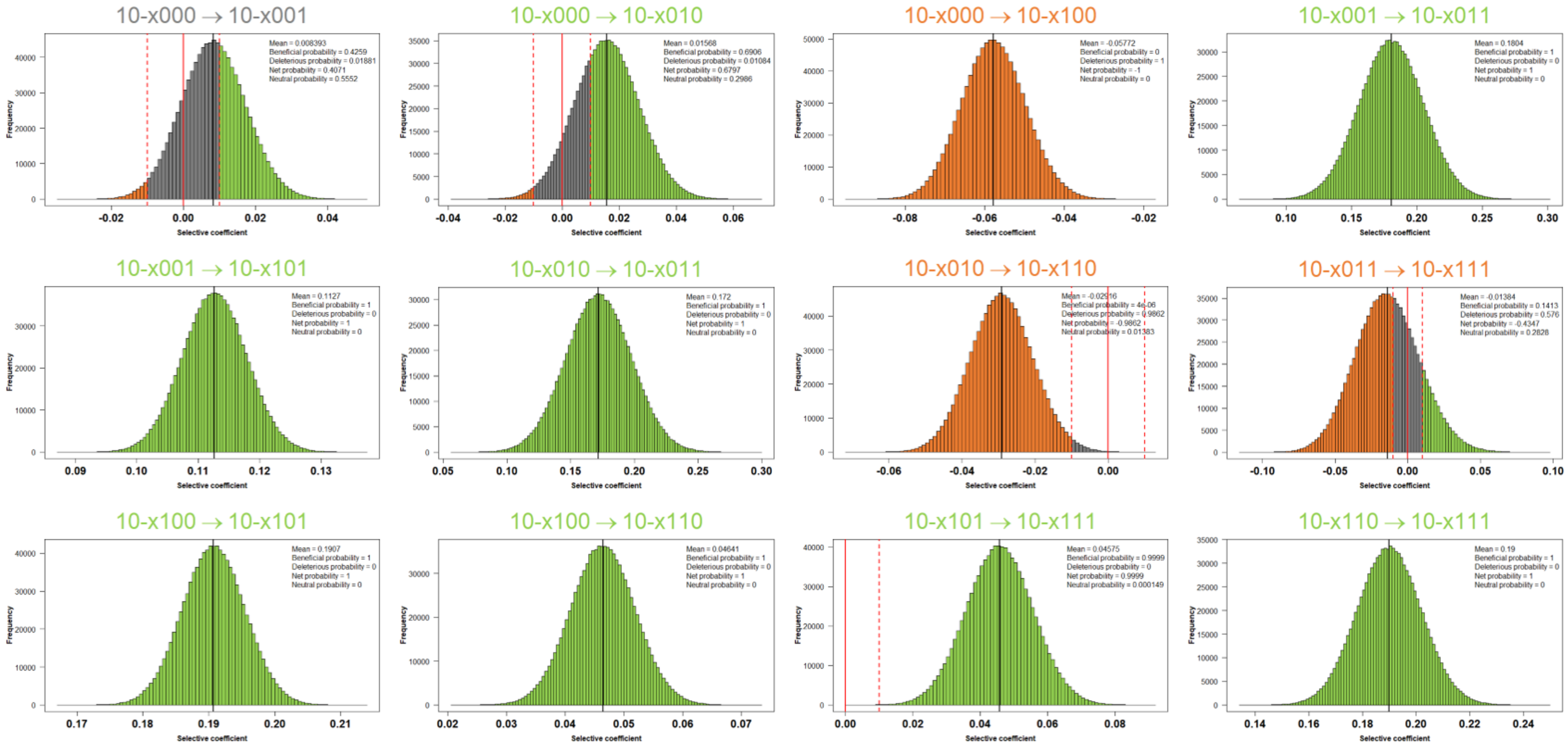

### Figure S7.

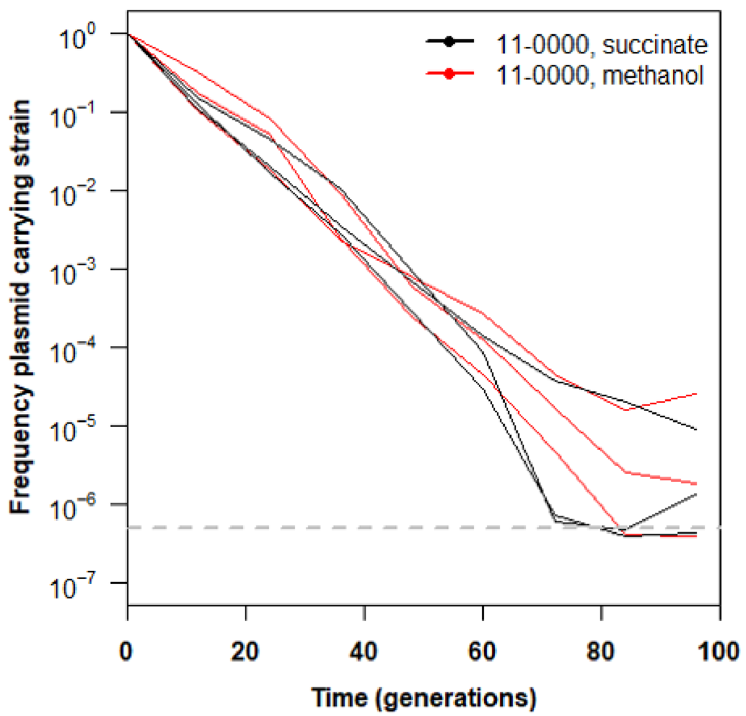
11-0000, succinate
11-0000, methanol
Frequency plasmid carrying strain
$10^{0}$
$10^{-1}$
$10^{-2}$
$10^{-3}$
$10^{-4}$
$10^{-5}$
$10^{-6}$
$10^{-7}$
0
20
40
60
80
100
Time (generations)

### Figure S8.

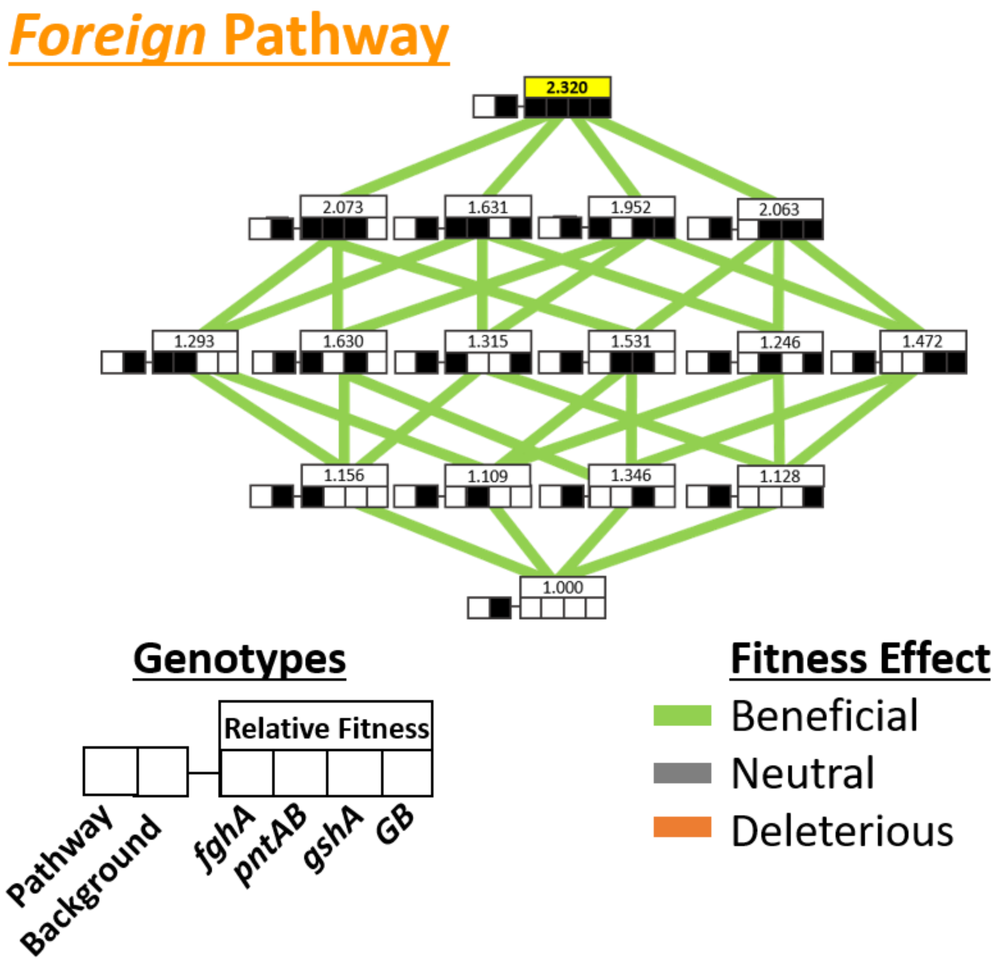
Foreign Pathway
2.320
2.073
1.631
1.952
2.063
1.293
1.630
1.315
1.531
1.246
1.472
1.156
1.109
1.346
1.128
1.000
Genotypes
Relative Fitness
Pathway
Background
fghA
pntAB
gshA
GB
Fitness Effect
Beneficial
Neutral
Deleterious
